# Knockout of the V-ATPase interacting protein Tldc2 in B-type kidney intercalated cells impairs urine alkalinization

**DOI:** 10.1101/2024.12.12.628211

**Authors:** Amity F. Eaton, Elizabeth C. Danielson, Leona J. Tu, Dennis Brown, Maria Merkulova

## Abstract

Intercalated cells (ICs) are acid-base regulatory cells in the kidney collecting duct that excrete either acid or base into the urine in response to systemic cues. A-ICs deliver protons into the tubule lumen via an apical proton pump (V-ATPase) and reabsorb base (bicarbonate) using the AE1 anion exchanger. B-ICs function in the opposite direction. They have basolateral V-ATPase and secrete bicarbonate into the lumen via the anion exchange protein, pendrin. The function of a third IC subtype, the non-A non-B IC which has apical pendrin and apical V-ATPase, is less well understood. We previously reported that members of the TLDc protein family interact with the V-ATPase and may regulate its function. TLDc proteins exhibit a distinct expression pattern in the kidney with RNAseq showing high, differential expression of Tldc2 in B-ICs. Here, we show by RNAscope imaging that Tldc2 is indeed expressed in B-ICs, but also in some non-A, non-B ICs. Using *Tldc2* knockout (*Tldc2*^-/-^) mice, we found that male and females had significantly lower urine pH than wild-type littermates, and their ability to increase urine pH in response to a bicarbonate load was impaired. In addition, *Tldc2*^-/-^ males developed hyperbicarbonatemia. *Tldc2*^-/-^ kidneys contained fewer B-ICs than wild-type mice, but more non-A, non-B ICs; therefore their sum and the number of A-ICs were unchanged. Furthermore, there was decreased basolateral accumulation of V-ATPase in *Tldc2*^/-/-^ B-ICs. These findings suggest that *Tldc2* is a novel gene involved in renal acid-base regulation and in addition, may serve as a differentiation marker for B-ICs.

**NEW & NOTEWORTHY:** Acid-base balance in the body is constantly changing but must be tightly controlled to be compatible with life. The kidney contains specialized cells that can excrete excess acid or base (bicarbonate) into the urine to maintain normal blood pH. The key protein involved in this process is called the V-ATPase. Here, we report that a novel V-ATPase interacting protein Tldc2 is critical for kidney bicarbonate secretion and is, therefore, a previously unrecognized acid-base regulatory gene.

## INTRODUCTION

The V-ATPase, or H^+^ATPase, is a proton pumping complex that is highly expressed in specialized cells in the kidney collecting duct, known as intercalated cells (ICs) (1). In these cells the V-ATPase complex contains several kidney-specific subunit isoforms and is trafficked to the plasma membrane, as opposed to more ubiquitous V-ATPases that acidify intracellular vesicles and organelles. The major function of ICs is to maintain acid-base homeostasis by adjusting the level and directionality of V-ATPase-dependent proton secretion by the kidney. There are two major types of ICs in the kidney, A- and B-type (2). A-type intercalated cells (A-ICs) express the V-ATPase at the apical membrane and the Cl^-^/HCO_3_ ^-^ anion exchanger 1 (AE1, also known as Slc4a1) at the basolateral membrane (3). Thus, A-ICs secrete protons into the urine via the V-ATPase and reclaim bicarbonate via AE1. Alternatively, B-type intercalated cells (B-ICs) express the V-ATPase at the basolateral membrane and the Cl^-^/HCO_3_ ^-^ anion exchanger pendrin (also known as Slc26a4) at the apical membrane (4–7). Thus, B-ICs secrete bicarbonate into the urine via pendrin and reclaim protons via the basolateral V-ATPase. To complicate matters, some ICs express pendrin at the apical membrane and V-ATPase either at the apical membrane or intracellularly – these are often referred to as non-A, non-B ICs (8). However, it is not yet clear if these cells are functionally distinct or a partially differentiated cell type in the process of transitioning to or from the A- or B-IC phenotype (8,9).

The V-ATPase is a large protein complex, consisting of a cytosolic V_1_ sector and a transmembrane V_0_ sector, each of which is comprised of multiple different subunits (1). There are two major mechanisms of V-ATPase regulation – via V_1_/V_0_ assembly/disassembly and via trafficking to specific membranes, such as the plasma membrane in kidney intercalated cells and in other proton secreting cell types (1). It is known that many proteins bind to the V-ATPase specifically and can directly regulate V-ATPase activity by affecting its trafficking and/or its assembly or disassembly (10). We found recently that all members of the so-called TLDc protein family interact with the V-ATPase (11), and it was shown recently that 4 out of 5 TLDc proteins not only bind to V-ATPase in vitro, but also induce V_1_ and V_0_ disassembly and consequently inhibit V-ATPase activity (12). Moreover, this process is conserved, since the yeast TLDc proteins Oxr1 and Rtc5 also inhibit V-ATPase activity by promoting its disassembly (13,14). Thus, the molecular function of TLDc proteins as endogenous disassembly factors and inhibitors of the V-ATPase is being uncovered.

Tldc2 is one member of this family of proteins, which all contain the highly conserved Tre2/Bub2/Cdc16 (TBC), Lysin Motif (LysM), Domain Catalytic (TLDc) hence their name - TLDc proteins. They are present in all eukaryotes from yeast to mammals and are mostly known for their role in protection against oxidative stress (15,16). In mammals there are five TLDc proteins, Oxr1, Ncoa7, Tbc1d24, Tldc1 and Tldc2 (17). Mutations in some TLDc protein genes result in various neurological disorders in humans (18–20) but the *in vivo* function of the family members Tldc1 and Tldc2 has not been studied yet.

Tldc2 is the smallest protein in the TLDc family and consists almost entirely of the conserved TLDc domain, which we showed is required for interaction with the V-ATPase (11). Similar to the other TLDc proteins, Tldc2 interacts with the V-ATPase and induces its disassembly *in vitro* (12). In a collaborative study, we previously found that in the kidney Tldc2 is highly expressed in pendrin-positive B-ICs and is among the top 25 B-IC selective transcripts, whereas it is not detectable in acid-secreting A-ICs (21). Importantly, a recent report highlighted the role of the V-ATPase in bicarbonate secretion by B-ICs during the renal defense against alkalosis (22). Based on these observations, the current study was carried out to test the hypothesis that Tldc2 is involved in regulation of V-ATPase activity in bicarbonate-secreting B-ICs and participates in homeostatic protection against systemic alkali loading.

## MATERIALS AND METHODS

### Mice

All animal studies were approved by the Massachusetts General Hospital Subcommittee on Research Animal Care, in accordance with the NIH, Department of Agriculture, and Association for the Assessment and Accreditation of Laboratory Animal Care requirements. Mice were housed in the Specific Pathogen-Free room at the Massachusetts General Hospital Animal Research Facility. Both male and female mice were used in the experiments. For all experiments, control mice were either littermates or age-and sex-matched mice from the same colony. The experimental unit was always a single mouse.

C57BL/6NJ-*Tldc2^em1(IMPC)J^*/Mmjax mouse strain (Mutant Mouse Resource & Research Centers (MMRRC) stock #42309) was generated by the Knockout Mouse Phenotyping Program (KOMP2) at The Jackson Laboratory (JAX, Bar Harbor, ME) using CRISPR/Cas9 technology. The details can be found at the JAX website: https://www.jax.org/strain/030887. These mice, hereafter called *Tldc2* knockout mice, carry a deletion of *Tldc2* exon 5 and 396 bp of flanking intronic sequence (including the splice acceptor and donor) that is predicted to cause a change of Tldc2 amino acid (aa) sequence after residue 143 and early truncation 24 aa later. Thus, the sequence VFKWTGHNSFFVKGDLDSLMMGSGSGQFGLWLDGDLYHGGSYPCATFNNEVLARREQFCIKELEAWVLS of the last 69 aa of the 212 aa -long Tldc2 protein, that is the third of its TLDc structural domain is replaced with the irrelevant sequence WPVWAVAGRRPVPWRKLPLCDLQQ. This substitution is predicted to result in a truncated unfolded protein that is non-functional.

One litter of heterozygous *Tldc2^+/-^* mice was received from JAX after its recovery from the cryo-archive and bred in-house to establish the *Tldc2* mouse colony. For genotyping, genomic DNA was extracted from mice tails using the KAPA Express Extract Kit (Kapa Biosystems, Wilmington, MA). Genotyping was performed by PCR using the KAPA Mouse Genotyping Kit (Kapa Biosystems) and F1/R1 primer pair (F1: 5’-TTGAAAGCAGGAAGCTTTGG-3’ and R1: 5’-ACTCTGGGACTTCCCCATCT-3’), that flanks Tldc2 exon 5 and detects both wild-type (+) and excised (-) alleles. To confirm that *Tldc2* full-length mRNA transcripts were absent from *Tldc2^-/-^* mice, end-point reverse transcription (RT-PCR) was carried out as briefly described. Total kidney RNA was isolated by using QIAshredder and RNeasy purification kit (Qiagen, Germantown, MD). RNA concentration was measured with a NanoDrop 2000 Spectrophotometer (Thermo Fisher Scientific, Waltham, MA). First-strand cDNA was then synthesized from 2 µg RNA using the High-Capacity RNA-to-cDNA Kit (Applied Biosystems, Waltham, MA) according to the manufacturer’s instructions. PCR reactions were performed using the Taq DNA Polymerase with Standard Taq Buffer (New England Biolabs, Ipswich, MA) on a T100 Thermal Cycler (Bio-Rad, Hercules, CA). The *Tldc2* exon 3-specific primer F2 5’-AAGGGATGGCTTCAGTCTGC-3’ and *Tldc2* mostly exon 5-specific primer R2 5’-CCACTTCCACTGCCCATCAT-3’ were used to detect *Tldc2* full-length transcripts. The F2/R2 RT-PCR product is detectable in *Tldc2^+/+^* mice and absent in *Tldc2^-/-^*mice. Beta actin (*Actb*)-specific primers 5’-TCGTGGGCCGCCCTAGGC-3’ and 5’-CAGCACAGCCTGGATGGC-3’ were used to amplify *Actb* cDNA as a loading and quality control.

### Blood gas, pH, electrolytes and urine pH measurements

In control experiments *Tldc2^-/-^* and *Tldc2^+/+^*(control) 4-5 month old mice were maintained on a standard rodent diet (Prolab® Isopro® RMH 3000, LabDiet, St. Louis, MO). Both male and female mice were used, and they had free access to water and food. For acid loading challenge, drinking water was substituted with 0.28 M NH_4_Cl/1% sucrose for 3 days. For alkali loading challenge, drinking water was substituted with 0.28 M NaHCO_3_/1% sucrose for 3 days. For measurement of blood gases, pH, sodium, chloride, free calcium and potassium, whole blood was collected from the submandibular vein of conscious mice directly into the BD Microtainer blood collection tube with heparin (cat # 365965, Becton Dickinson and Co, Franklin Lakes, NJ). It was then immediately tested using the SC80 ABL80 cassette (50-test, 30-days Full panel with QC3 (no Glu), cat # 945-670, Radiometer America, Westlake, OH) and the SP80 ABL80 Solution Pack (cat # 944-174, Radiometer America) on a Radiometer ABL80Flex analyzer (Radiometer America). Blood bicarbonate concentration was calculated from the measured pH and pCO_2_ values using the Henderson-Hasselbalch equation. For spot urine pH measurement, mouse urine was collected directly onto Hydrion Urine & Saliva pH Paper 5.5-8.0 (Micro Essential Laboratory Inc., Brooklyn, NY) or colorpHast pH 7.5-14.0 strips (MilliporeSigma, Burlington, MA) and pH was read immediately.

### Antibodies

Commercial primary antibodies used in this study were rabbit monoclonal anti-AE1 at 1:200 (for immunofluorescence (IF)) (cat # 20112S, Cell Signaling, Danvers, MA), rabbit polyclonal anti-pendrin at 1:100 (IF) (cat # PA5-42060, ThermoFisher Scientific, Waltham, MA) and β-Actin-HRP at 1:1000 (for Western blotting (WB)) (cat # 5125S, Cell Signaling, Danvers, MA). Commercial secondary antibodies used in this study for immunofluorescence were goat anti-chicken Alexa 555 (cat # A32932, 1:1000, Invitrogen, Waltham, MA), goat anti-chicken Alexa 647 (cat # A32933, 1:1000, Invitrogen, Waltham, MA), goat anti-rabbit Alexa 488 (cat # A32731, 1:1000, Invitrogen, Waltham, MA), and for western blotting goat anti-rabbit-HRP (cat # ab97080, 1:5000, Abcam, Waltham, MA) and donkey anti-chicken-HRP (cat # 703-035-155, Jackson ImmunoResearch, West Grove, PA). The chicken polyclonal anti-B1 V-ATPase (56-kDa subunit) antibody (1:200, IF and 1:1000, WB) was raised against a 13-amino acid peptide (QGAQQDPASDTAL), corresponding to the COOH-terminal sequence of mouse V-ATPase. This antibody was produced, affinity purified, and characterized previously in our laboratory (23). The non-commercial rabbit polyclonal anti-pendrin antibody was a kind gift from Dr. Susan Wall, Emory University and used at 1:1000 (WB) (7).

### Whole Body Perfusion Fixation and Tissue Preparation

Four- to five-month-old control (*Tldc2*^+/+^) and *Tldc2* knockout (*Tldc2*^-/-^) mice were anaesthetized by IP injection 50 mg/kg body with Nembutal (NDC: 11695-4862-5, Covetrus, Dublin, OH). Animals were allowed to reach proper anesthetic depth over a period of 5 min, which was confirmed by lack of a response to a toe pinch. Mice were then perfused through the left ventricle of the heart for 1-2 min at 17 mL/min with 37°C PBS. For western blotting and PCR, the renal artery feeding the right kidney was clamped with a hemostat and the right kidney was excised, capsule removed, rinsed with PBS, and flash frozen in liquid nitrogen and stored at −80°C until ready for lysate preparation (see below). For BaseScope and immunofluorescence assays, the left kidney was perfusion fixed *in situ* with 37°C 4% paraformaldehyde in PBS or paraformaldehyde-lysine-periodate fixative (PLP; 4% paraformaldehyde, 75 mM lysine-HCl, 10 mM sodium periodate, and 0.15 M sucrose, in 37.5 mM sodium phosphate), respectively, for 5 min at 17 mL/min. The kidney was extracted, sectioned into five circumferential pieces, and fixed by immersion in PFA or PLP overnight at 4°C with rocking. Tissues for BaseScope were sequentially immersed in 10, 20, and 30% sucrose in PBS at 4°C until the tissues sank and snap frozen in Tissue-Tek OCT compound (Sakura Finetek USA, Torrance, CA) in liquid nitrogen and stored at −80°C until ready for cryosectioning. Tissues for indirect immunofluorescence were rinsed 5×1 h at RT in PBS and stored at 4°C in PBS + 0.02% sodium azide until ready for cryoprotection in 30% sucrose in PBS for 72 h at 4°C.

Tissues prepared for immunofluorescence, as described above, were embedded in Tissue-Tek OCT compound on a specimen disk (Sakura Finetek USA, Torrance, CA), and frozen at −20°C. Tissues were sectioned at 5 μm on a Leica CM3050S cryostat (Leica Microsystems, Bannockburn, IL), collected onto Superfrost Plus microscope slides (Thermo Fisher Scientific, Rockford, IL), and stored at −80°C until ready for use.

### BaseScope fluorescent detection of in situ RNA

First, the validity of the BaseScope assay in murine kidney in our hands was established using the positive control probes (ppib) provided with the BaseScope^TM^ assay kit (Advanced Cell Diagnostics (ACD) Bio, Newark, CA) kit, and which should be expressed in every cell at moderate-to-high levels (Supplemental Fig. S1*A*). Then, negative control probes (dapb) were used, recognizing a bacterial protein that should not be expressed in mammalian cells at all (Supplemental Fig. S1*B*). After that, BaseScope assay was performed using a custom *Tldc2* probe (BA-Mm-Tldc2-1zz-st-C1, Cat No. 1333581-C1) designed by ACD Bio. The BaseScope^TM^ *Tldc2* probe targets a 482-526 nucleotide sequence of Tldc2 mRNA NM_001177439. BaseScope assay was performed following ACD Bio’s BaseScope RED protocol as adapted for fixed frozen tissues. Briefly, slides were baked for 30 min @ 60°C in a dry oven followed by post-fixation in 4% PFA in PBS for 15 min at 4°C, and sequential dehydration of slides in 50, 70, and 100% EtOH for 5 min at RT. Sections were pretreated with H_2_O_2_ (ACD Bio) for 10 min at RT and antigen retrieval was performed by boiling slides for 5 min in 1X Target Retrieval Reagent (ACD Bio), followed by incubation with Protease III (ACD Bio) for 30 min at 40°C prior to probe hybridization, amplification, and labeling with Fast-RED (ACD Bio). Indirect immunofluorescence labeling, as described below, was performed subsequently.

### Indirect Immunofluorescence Labeling and Confocal Image Capture

For each experiment, cryosections from at least 4 control and 4 *Tldc2^-/-^* mice were incubated in parallel. Sections were rehydrated for 3×5 min in PBS. For immunofluorescence with V-ATPase antibodies sections were incubated with 1% (wt/vol) SDS for 4 min for antigen retrieval (24). After antigen retrieval, sections were washed for 3×5 min in PBS and incubated for 30 min in 1% (wt/vol) bovine serum albumin (BSA) in PBS. The sections were incubated for 90 min at RT, or overnight at 4°C, with the primary antibody diluted in 1% (wt/vol) BSA in PBS. After primary antibody incubation sections were washed 3×5 min with PBS and the secondary antibody was applied for 1h at room temperature. Finally, the slides were rinsed again 3×5 min in PBS and mounted with SlowFade Diamond Antifade Mountant (ThermoFisher Scientific, Waltham, MA) with DAPI as a nuclear stain (Vector Laboratories, Burlingame, CA). Confocal images were acquired using an LSM 800 confocal laser scanning microscope (Carl Zeiss Microscopy, Thornwood, NY), controlled by ZEN 2 (blue edition) software (Carl Zeiss Microscopy). For comparative semiquantitative analysis (see below), confocal images of control and *Tldc2^-/-^* kidneys were acquired under identical conditions and imaging parameters. Images were postprocessed using Adobe Photoshop CS4 image-editing software (Adobe Systems, San Jose, CA).

### Fluorescence Image Analysis

A-type ICs were identified by their expression of the B1 V-ATPase subunit at the apical membrane and anion exchanger 1 (AE1) on the basolateral membrane. Non-A Non-B type ICs were defined by their localization to the cortex and apical B1 V-ATPase and pendrin expression. B-type ICs were identified by their localization in the cortex and basolateral/diffuse expression of B1 V-ATPase and apical expression of pendrin. Cells for which the nucleus or the apical membrane were not clearly visible due to the orientation of the cut were excluded from analysis. Images from four animals per genotype, and two per sex, were quantified for each parameter. Images were analyzed using ImageJ version 1.53a (NIH, Bethesda, MD) and ZenBlue software (Carl Zeiss Microscopy, White Plains, NY), and data were imported into Microsoft Excel version 16 (Microsoft, Redmond, WA), and GraphPad Prism version 10 (GraphPad Software, San Diego, CA) was used for further graphical and statistical analysis.

### V-ATPase Apical/Basolateral Domain Fluorescence Intensity

Apical and basolateral domain fluorescence intensity of the B1 V-ATPase and apical domain fluorescence intensity of pendrin were quantified separately. Briefly, using the ImageJ “line tool”, a line of 0.5 μm width was drawn across each cell starting from, and perpendicular to, the apical membrane and adjacent to the nucleus. From this line, a “line intensity profile” was generated in ImageJ, which plots fluorescence intensity on the y-axis and distance from the apical membrane in microns on the x-axis. The mean fluorescence intensity of a circular ROI drawn adjacent to the apical membrane of the cell was subtracted from intensity values as background. The estimated area under the curve for each 0.05 μm rectangle was calculated using the equation:

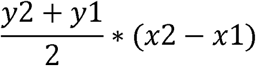

The sum of all the estimated areas under the curve was considered the total area under the curve. The apical domain was defined as the sum of the estimated areas under the curve from 0 to 2 μm and the basolateral domain was defined as the sum of the estimated areas under the curve 2 μm from the ultimate value. Apical or basolateral domain fluorescence intensity was calculated as the percentage of apical or basolateral domain fluorescence intensity relative to the total cell fluorescence intensity. The percentage of apical or basolateral domain fluorescence intensity for 25 cells was averaged per animal.

### Quantification of percentage of pendrin-positive cells (non-A ICs) relative to total ICs

2 mm x 2 mm mosaic images from the cortex of kidney cryosections were captured where at least 100 total ICs were quantified. The number of pendrin-negative ICs (A-ICs) and pendrin-positive ICs (non-A ICs) was quantified and the percentage of pendrin-positive ICs relative to total ICs was calculated.

### Semiquantitative Western blotting

Mouse kidneys were dissected, and transverse sections from the central region of each mouse kidney were lysed in ice-cold RIPA buffer (50 mM Tris·HCl, 150 mM NaCl, 1% NP-40, 0.5% sodium deoxycholate, and 0.1% SDS, pH 7.4; Boston BioProducts, Ashland, MA,), containing Complete Protease Inhibitor cocktail (cat # 11697498001, Roche Applied Science, Indianapolis), at 1 ml of lysis buffer per 100 mg of kidney tissue. Tissues were homogenized with a motorized tissue grinder (cat # 12-1413-61, Fisher Scientific), allowed to lyse for 20 min on ice and clarified via centrifugation at 16,000 x *g* for 20 min at 4°C. Total protein concentration was determined using a Pierce BCA Protein Assay Kit (Thermo Fisher Scientific, Waltham, MA). An equal amount of each protein sample was loaded on Novex NuPAGE 4–12% Bis-Tris gels (Thermo Fisher Scientific). Protein separation, transfer to PVDF membrane (Thermo Fisher Scientific) and western blotting were performed conventionally. Equal loading and protein transfer to the membrane were confirmed by Ponceau S staining (Sigma-Aldrich, St. Louis, MO). The chemiluminescence images were digitally captured using G:BOX mini imaging system (Syngene, Synoptics) and quantified using ImageJ software [version 2.3.0/1.53q; National Institutes of Health] (25).

### Statistical Analysis

Statistical significance was determined using a two-tailed unpaired t-test using Prism 9 software (GraphPad Software, San Diego, CA). A P-value ≤.05 was considered statistically significant. Graphs were plotted with Prism 9 software. Experimental values are reported as means ±standard error of the mean (SEM).

## RESULTS

### Tldc2 is specifically expressed in B-intercalated cells and a subset of non-A non-B-intercalated cells in the mouse kidney

In high-throughput single cell RNA transcriptomics studies we, and others, found that *Tldc2* was one of the most highly differentially expressed genes in B-ICs in mouse kidney (21). To confirm this data and study where Tldc2 is expressed *in situ* we performed fluorescence microscopy imaging using kidney tissue cryosections. Since we were unable to find reliable Tldc2 antibodies, we used BaseScope, a version of the RNAscope technique, to identify where the *Tldc2* RNA was expressed in the mouse kidney. Positive BaseScope “staining” appears in the form of multiple fluorescent puncta, which corresponds to the location of RNA transcripts for the protein of interest. After confirmation of the validity of the BaseScope assay in murine kidney in our hands, as described in MATERIALS AND METHODS, we used specific probes designed against Tldc2 RNA, together with co-staining of the same sections with anti-pendrin and anti-B1 antibodies to identify and distinguish different types of ICs. We found that Tldc2 RNA was present in pendrin-positive, basolateral B1-positive B-ICs (arrows), but not in pendrin-negative, apical B1-positive A-ICs (arrowheads) (Fig. 1*A*). Intriguingly, we saw that in addition Tldc2 puncta were detected in a small subset of pendrin-positive non-A non-B ICs, that express B1 either apically or intracellularly (Fig. 2). Thus, these data confirm the expression of *Tldc2* mRNA in B-ICs as seen by single cell RNAseq and extend this finding to include a small population of non-A, non-B ICs.

**Figure 1.**
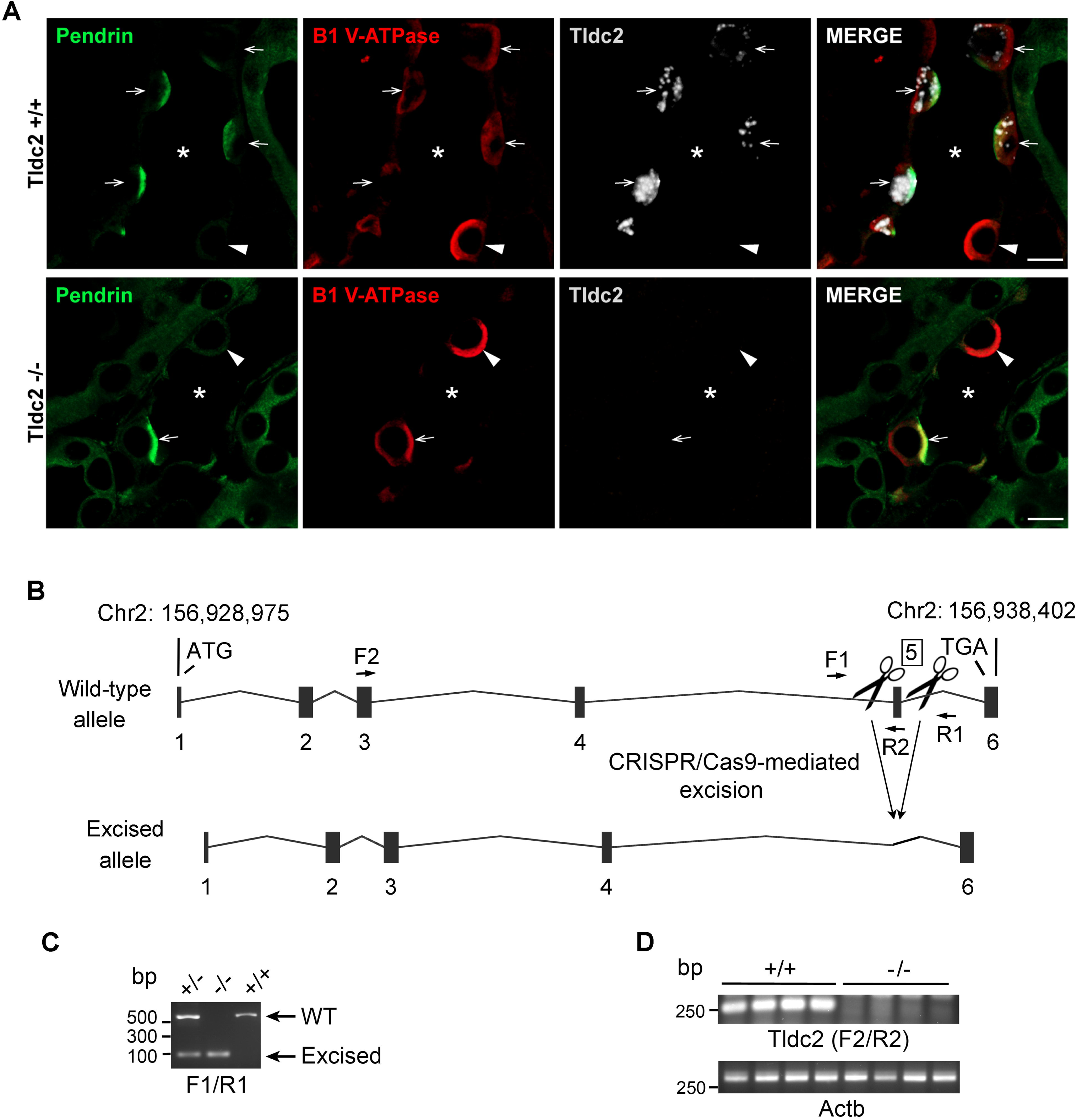
Tldc2 is specifically expressed in pendrin-positive intercalated cells from Tldc2^+/+^ mice and is absent in kidneys from Tldc2^-/-^ mice. (A) Immunofluorescence images taken from the cortex of kidneys from wildtype Tldc2^+/+^ (top) and knockout Tldc2^-/-^ (bottom) mice after performing RNA scope using specific probes designed against Tldc2 (spots, white) and co-staining with specific antibodies against pendrin (green) and the B1 V-ATPase (red). A positive RNA scope signal appears as a single intracellular punctum (white). Apical pendrin expression with basolateral B1 expression was used to identify B-ICs (arrows) and apical B1 expression and the absence of pendrin expression was used to identify A-ICs (arrowheads). Tubule lumen is indicated with an asterisk (*). Scale bars = 10 µM. (B) Schematic representation of *Tldc2* gene and its wild-type (WT) and excised alleles. *Tldc2* gene is located at mouse Chromosome 2: 156,928,975-156,938,402 forward strand (Genome Build GRCm39:CM000995.3) and contains 6 exons. Excised allele was generated by CRISPR/Cas9-mediated deletion of exon 5 and flanking sequences as described in Materials and Methods. Positions of forward and reverse primer pairs used for mouse genotyping (F1/R1) and end-point RT-PCR (F2/R2) are indicated. (C) Representative results of PCR genotyping of genomic DNA extracted from tails of wild-type (+/+), heterozygous (+/-) and homozygous (-/-) *Tldc2* knockout mice. Primer pair F1/R1 detects both WT (+) and excised (-) alleles and amplifies a 562 bp WT fragment and a 92 bp excised fragment. The inferred genotypes are shown above. (D) The absence of *Tldc2* full-length transcripts in *Tldc2^-/-^* mice was confirmed by end-point RT-PCR using F2 and R2 primers, specific for *Tldc2* exons 2 and 5 respectively. Total RNA was isolated from 4 WT (+/+) and 4 homozygous (-/-) *Tldc2* knockout mouse kidneys, reverse transcribed and amplified by PCR. Primer pair F2/R2 amplified a 264 bp fragment within WT (+/+) *Tldc2* RNA transcript only. *Actb*-specific primer pair was used as a PCR loading control and amplified a 302 bp RNA fragment from both *Tldc2^+/+^* and *Tldc2^-/-^* mice, as expected.

**Figure 2.**
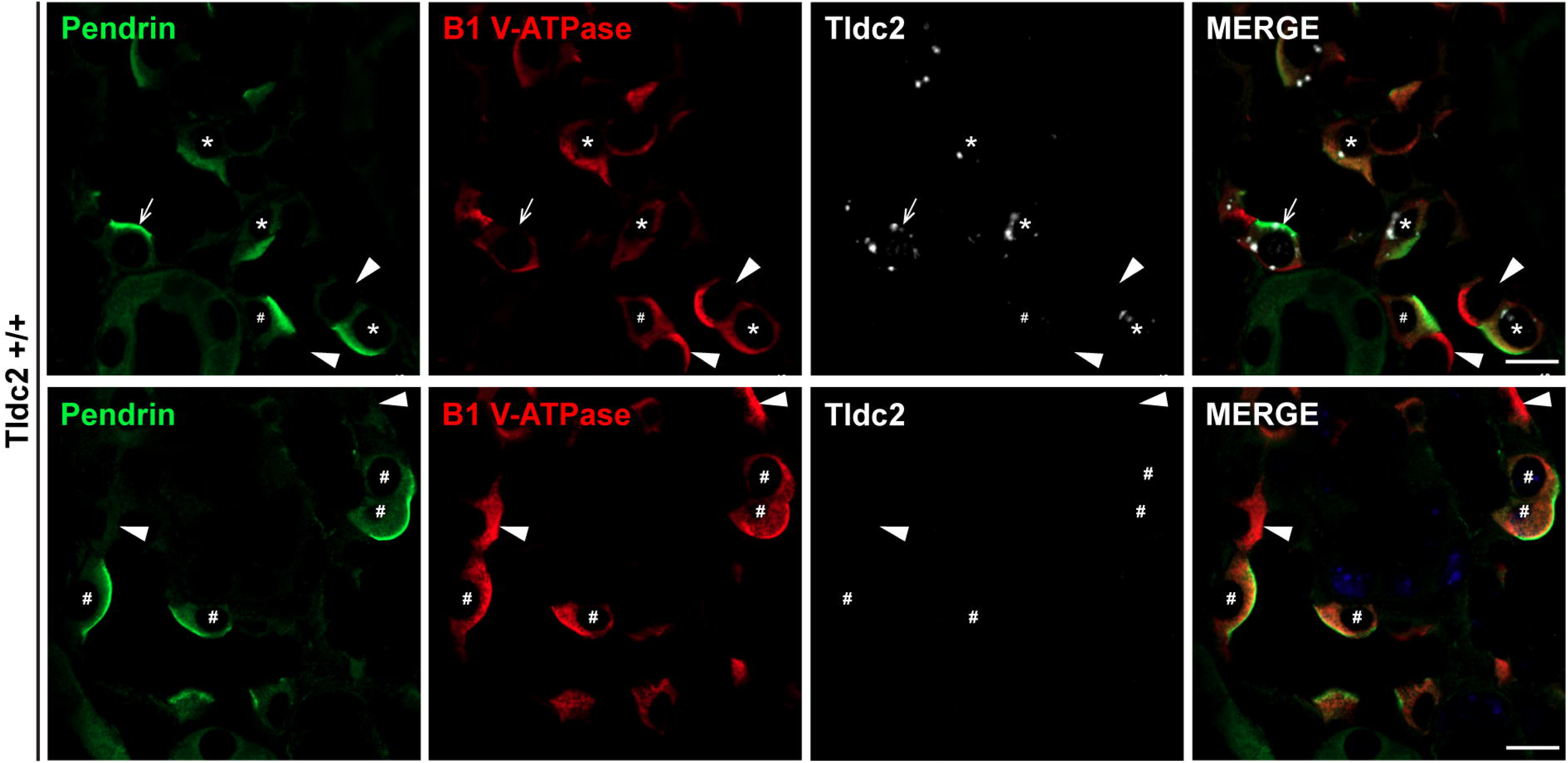
Tldc2 is specifically expressed in B-type intercalated cells (B-ICs) and a subset of non-A non-B-type intercalated cells (non-A non-B ICs) from Tldc2^+/+^ mice. Immunofluorescence images taken from the cortex of kidneys from wildtype Tldc2^+/+^ mice after performing RNA scope using specific probes designed against Tldc2 (spots, white) and co-staining with specific antibodies against pendrin (green) and the B1 subunit of the V-ATPase (red). A positive RNA scope signal appears as a single intracellular punctum (white). Apical pendrin expression with basolateral B1 expression was used to identify B-ICs, apical pendrin expression with apical B1 expression was used to identify non-A non-B ICs, and apical B1 expression and the absence of pendrin expression was used to identify A-ICs. Arrows indicate B-ICs, asterisks (*) indicate Tldc2-positive non-A non-B ICs, pound signs (#) indicate Tldc2-negative non-A non-B ICs, arrowheads indicate A-ICs. Scale bars = 10 µM.

### Confirmation of Tldc2 loss in the kidneys of Tldc2 knockout mice, generated by the high-throughput Knockout Mouse Phenotyping Program

The Tldc2 global knockout mouse strain was generated at The Jackson Laboratory as part of the Knockout Mouse Phenotyping Program (KOMP2). The gene targeting strategy is shown in Fig. 1*B* and more details are included in the MATERIALS AND METHODS section. Briefly, Cas9-mediated excision resulted in the deletion of exon 5, frame-shifted after exon 4, resulting in production of a truncated, non-functional protein. We received a litter of heterozygous *Tldc2^+/-^*mice, which was recovered from a cryo-archive at The Jackson Laboratory. We confirmed the *Tldc2^+/-^* genotype by PCR genotyping on tail genomic DNA using primer pair F1/R1, flanking exon 5 outside the deletion and detecting both wild-type and excised alleles (Fig. 1*C*). These *Tldc2^+/-^*mice were then intercrossed to generate littermate *Tldc2*^+/+^, *Tldc2*^+/-^ and *Tldc2*^-/-^ mice and establish a *Tldc2* global knockout mouse colony in-house. After genotyping, absence of full-length *Tldc2* transcripts in *Tldc2*^-/-^ mice was validated by end-point RT-PCR using total kidney RNA and primer pair F2/R2 (Fig. 1*D*). We further confirmed the knockout of *Tldc2* at the RNA level in the kidneys of our *Tldc2^-/-^* mice by BaseScope. Whereas there were numerous fluorescent puncta present in the B-IC from the *Tldc2*^+/+^ kidneys (Fig. 1*A*, upper panel), no fluorescent puncta were detectable in our *Tldc2*^-/-^ mice, confirming that Tldc2 is knocked out in the kidneys of these animals (Fig. 1*A*, lower panel).

### Tldc2^-/-^ knockouts have persistently low urine pH and male, but not female, Tldc2^-/-^ mice develop hyperbicarbonatemia without alkalemia under standard and alkali load diet

Overall, anatomically *Tldc2*^-/-^ mice were indistinguishable from *Tldc2*^+/+^ controls and looked normal without gross physiological abnormalities. To study if there is any acid-base imbalance in *Tldc2*^-/-^ mice on the standard diet we first measured their urine pH and found that it was significantly lower in *Tldc2*^-/-^ than in *Tldc2*^+/+^ control mice (5.78±0.03 vs. 6.13±0.05 in males, Fig. 3 and Table 1; and 5.88±0.04 vs. 6.17±0.08 in females; Fig. 4 and Table 1). We then measured blood pH, HCO3- and pCO2 levels in both *Tldc2*^-/-^ males and females on the standard diet. We found that *Tldc2*^-/-^ male, but not female, mice developed a mild hyperbicarbonatemia in comparison with the corresponding *Tldc2*^+/+^ controls (22.60±0.69 vs. 20.38±0.70 mM in males, Fig. 3 and Table 1; and 22.38±0.98 vs. 22.57±1.19 in females; Fig. 4 and Table 1). However, this hyperbicarbonatemia was accompanied by hypercapnia (44.51±0.87 vs. 39.20±1.70 mmHg, Fig. 3 and Table 1) and, therefore, blood pH in *Tldc2*^-/-^ males was not significantly different from *Tldc2*^+/+^ controls (7.33±0.01 vs. 7.34±0.01, Fig. 3 and Table 1) due to respiratory compensation. Blood pH in *Tldc2*^-/-^ females was also normal compared to *Tldc2*^+/+^ females (7.35±0.02 vs. 7.34±0.02, Fig. 4 and Table 1). Therefore, both *Tldc2*^-/-^ males and females were not alkalemic despite aciduria.

**Figure 3.**
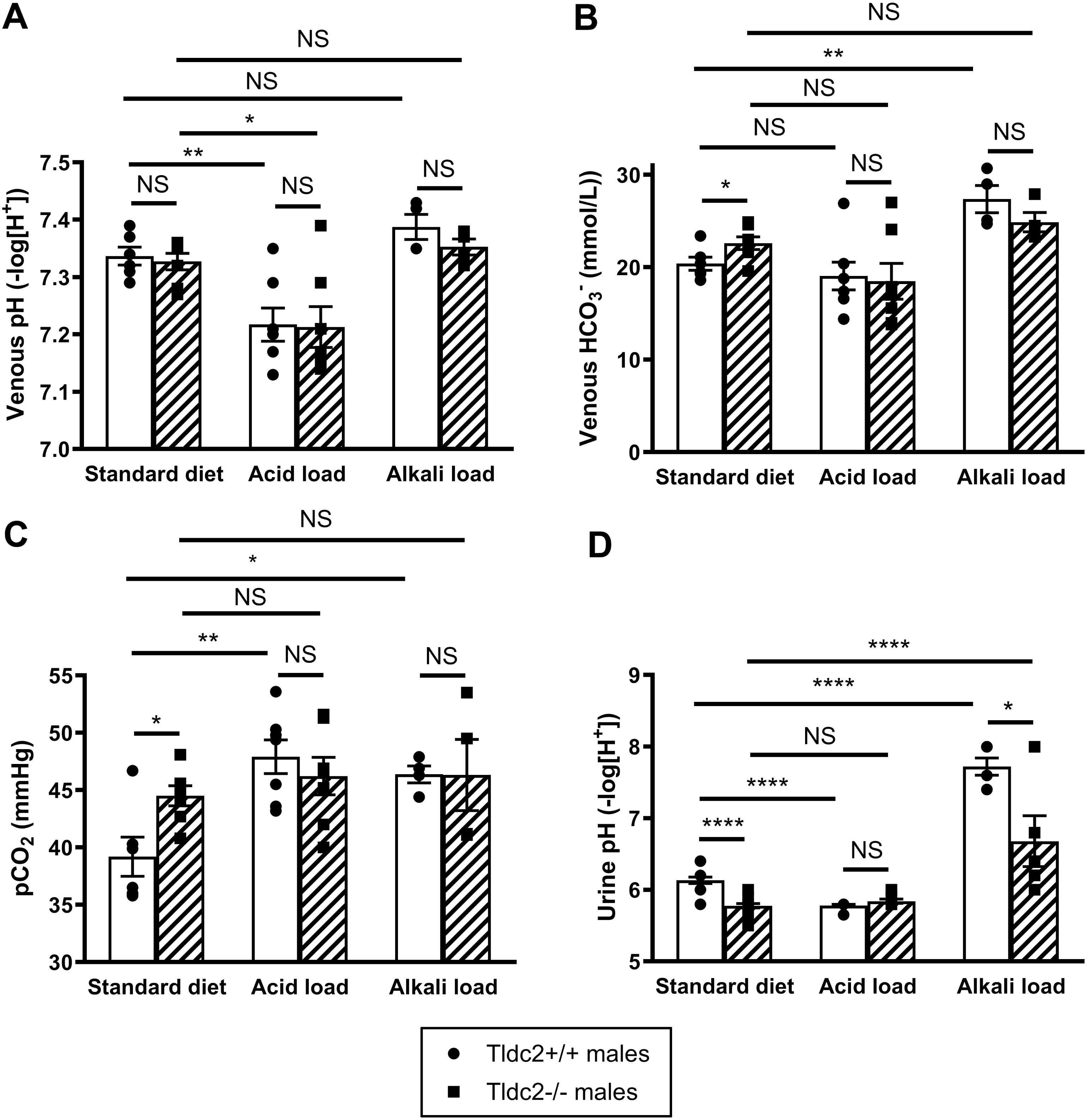
*Tldc2^-/-^*male mice demonstrate hyperbicarbonatemia compensated by hypercapnia and have persistently low urine pH. (A) Blood pH in *Tldc2^-/-^* male mice is not significantly different from *Tldc2^+/+^* control males on a standard rodent diet. Both *Tldc2^-/-^* and *Tldc2^+/+^* male mice developed acidemia after acid loading with 0.28 M NH_4_Cl for 3 days, without significant difference between genotypes. Blood pH did not change significantly after alkali loading with 0.28 M NaHCO_3_ for 3 days in ether *Tldc2^-/-^* or *Tldc2^+/+^* males. (B) Blood HCO ^-^ levels were significantly higher in *Tldc2^-/-^* than in *Tldc2^+/+^*males maintained on a standard diet. After acid loading, blood HCO ^-^ levels did not change significantly in either *Tldc2^-/-^* or *Tldc2^+/+^*males. After alkali loading, blood HCO ^-^ levels were increased in *Tldc2^+/+^*, but not in *Tldc2^-/-^* males. (C) Blood partial pressure of carbon dioxide (pCO_2_) was significantly higher in *Tldc2^-/-^* than in *Tldc2^+/+^*males maintained on a standard diet. After acid loading, blood pCO_2_ levels were significantly increased in *Tldc2^+/+^*, but not in *Tldc2^-/-^* males. After alkali loading, blood pCO_2_ levels were also significantly increased only in *Tldc2^+/+^*. (D) Urine pH was significantly lower in *Tldc2^-/-^* males compared to *Tldc2^+/+^* controls on a standard diet. Acid loading resulted in a more acidic urine pH in *Tldc2^+/+^* control males, but not in *Tldc2^-/-^* males, whose urine pH was very low even on a non-acidified diet. After alkali loading urine pH was significantly increased in male mice of both genotypes, but it remained significantly lower in *Tldc2^-/-^*than in *Tldc2^+/+^* males. All values are means ± SEM. * is for p-value < 0.05, ** is for p-value < 0.01, **** is for p-value < 0.0001; t-test. NS = non-significant. The actual p-values and other details can be found in Tables 1 and 2.

**Figure 4.**
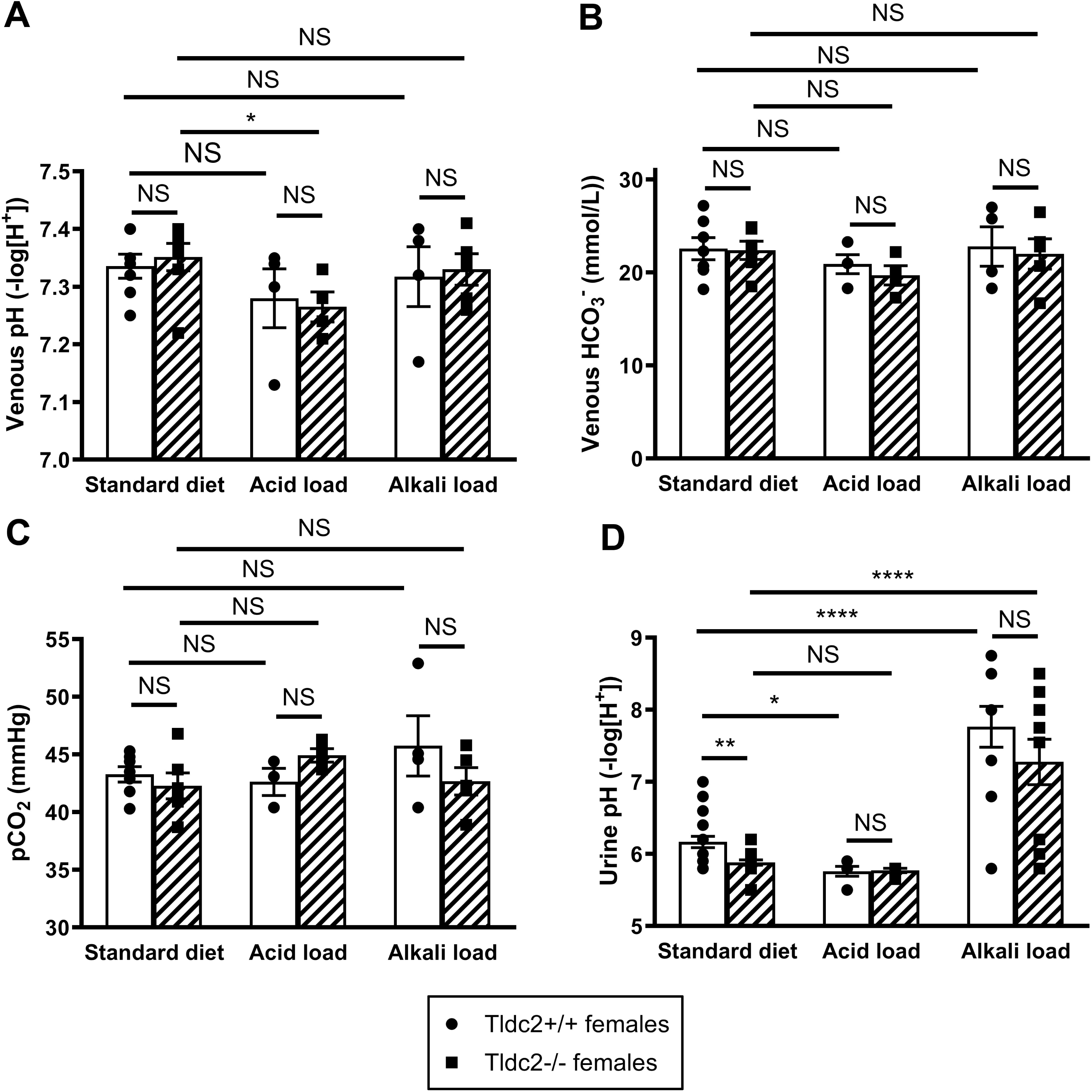
*Tldc2^-/-^* female mice demonstrate persistently low urine pH, but no significant changes in blood pH, bicarbonate or carbon dioxide levels. (A) Blood pH in *Tldc2^-/-^* female mice is not significantly different from *Tldc2^+/+^* control females on a standard rodent diet, after acid loading with 0.28 M NH_4_Cl for 3 days or after alkali loading with 0.28 M NaHCO_3_ for 3 days. Blood pH was significantly reduced after acid loading in *Tldc2^-/-^* females only. Blood pH did not change significantly after alkali loading in ether *Tldc2^+/+^* or *Tldc2^-/-^*females. (B) and (C) Blood HCO ^-^ concentration and blood partial pressure of carbon dioxide in *Tldc2^-/-^*female mice are not significantly different from *Tldc2^+/+^* control females maintained on a standard diet. They did not change significantly after acid loading with 0.28 M NH_4_Cl for 3 days or after alkali loading with 0.28 M NaHCO_3_ for 3 days in ether *Tldc2^+/+^*or *Tldc2^-/-^* females and there were no statistically significant differences between genotypes. (D) Urine pH was significantly lower in *Tldc2^-/-^* females compared to *Tldc2^+/+^* controls on a standard diet. Acid loading resulted in a more acidic urine pH in *Tldc2^+/+^*control females, but not in *Tldc2^-/-^* females, whose urine pH was very low even on a non-acidified diet. After alkali loading urine pH was significantly increased in both *Tldc2^+/+^* and *Tldc2^-/-^*females and there was no statistically significant difference between genotypes due to high variability of response of female mice to bicarbonate loading. All values are means ± SEM. * is for p-value < 0.05, ** is for p-value < 0.01, **** is for p-value < 0.0001; t-test. NS = non-significant. The actual p-values and other details can be found in Tables 1 and 2.

**Table 1.**
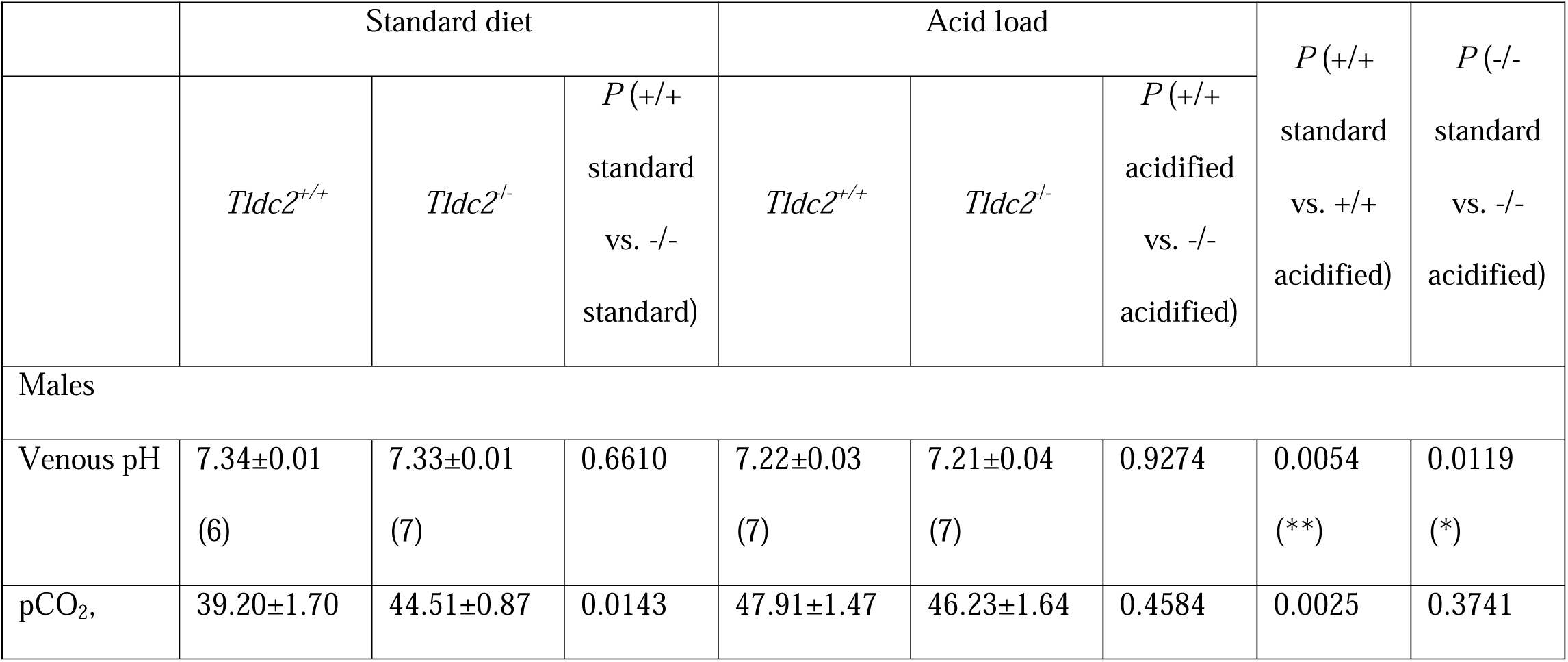
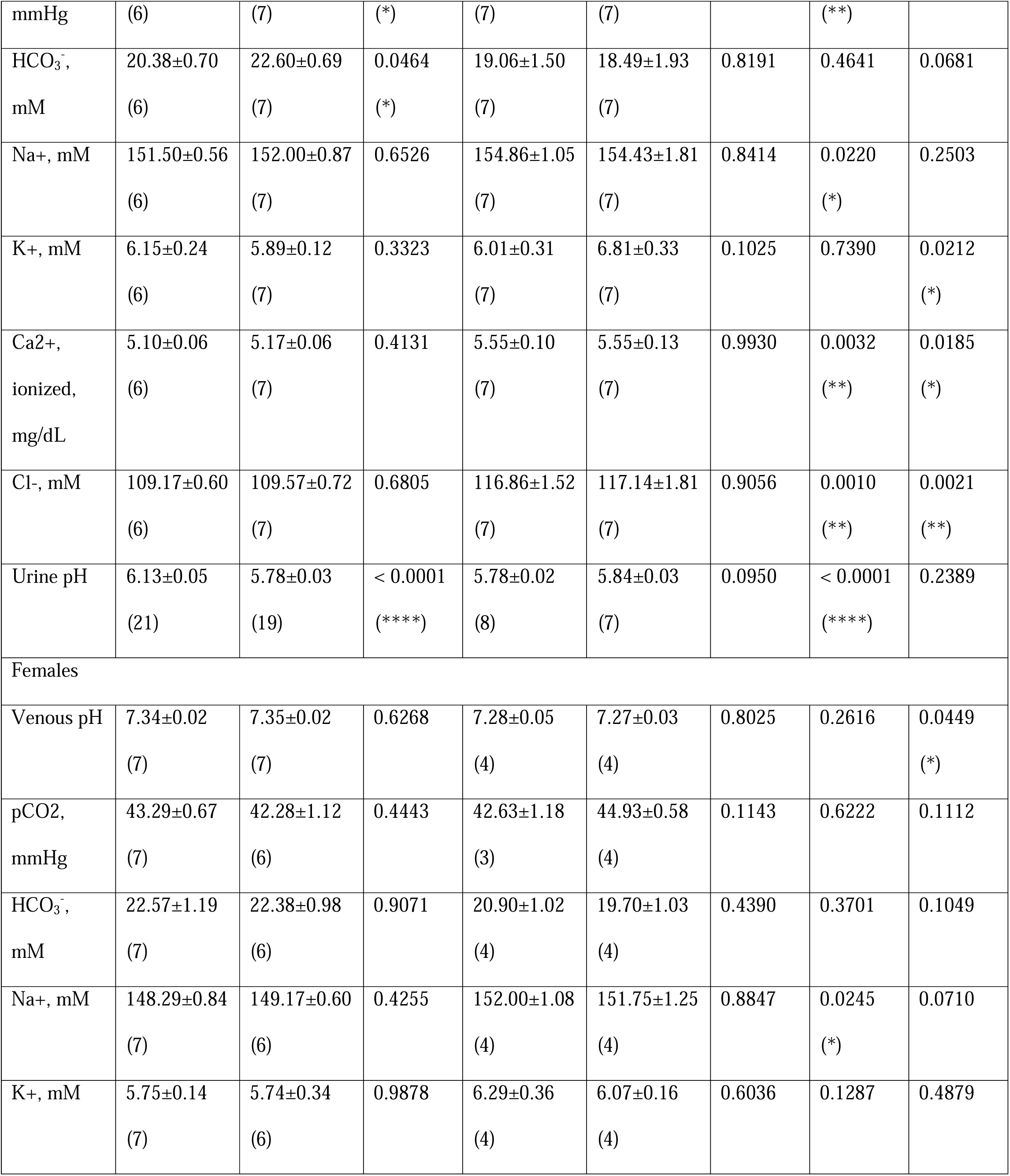
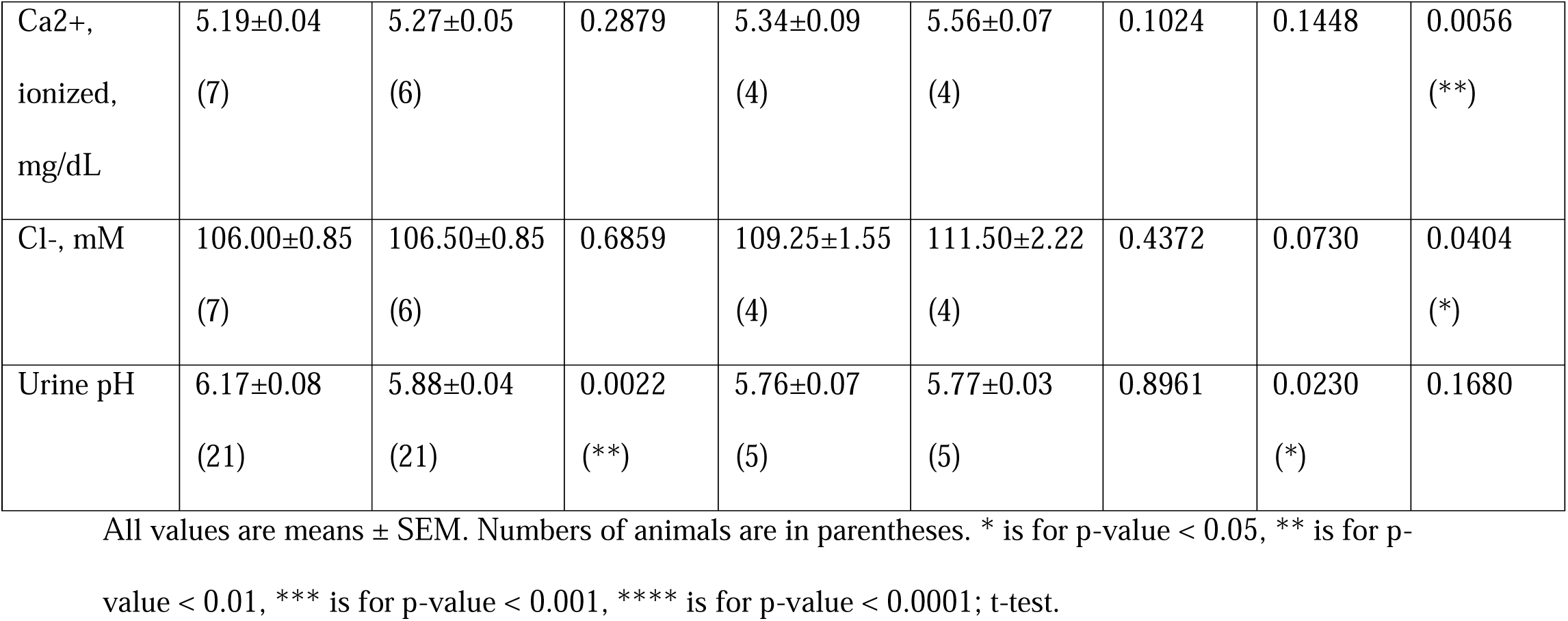
Summary of NH_4_Cl –loading of male and female mice. Venous blood gases, pH and major electrolytes, and urine pH of *Tldc2*^-/-^ knockout and *Tldc2^+/+^* control mice on a standard rodent diet and after acid loading with 0.28 M NH_4_Cl for 3 days.

**Table 2.**
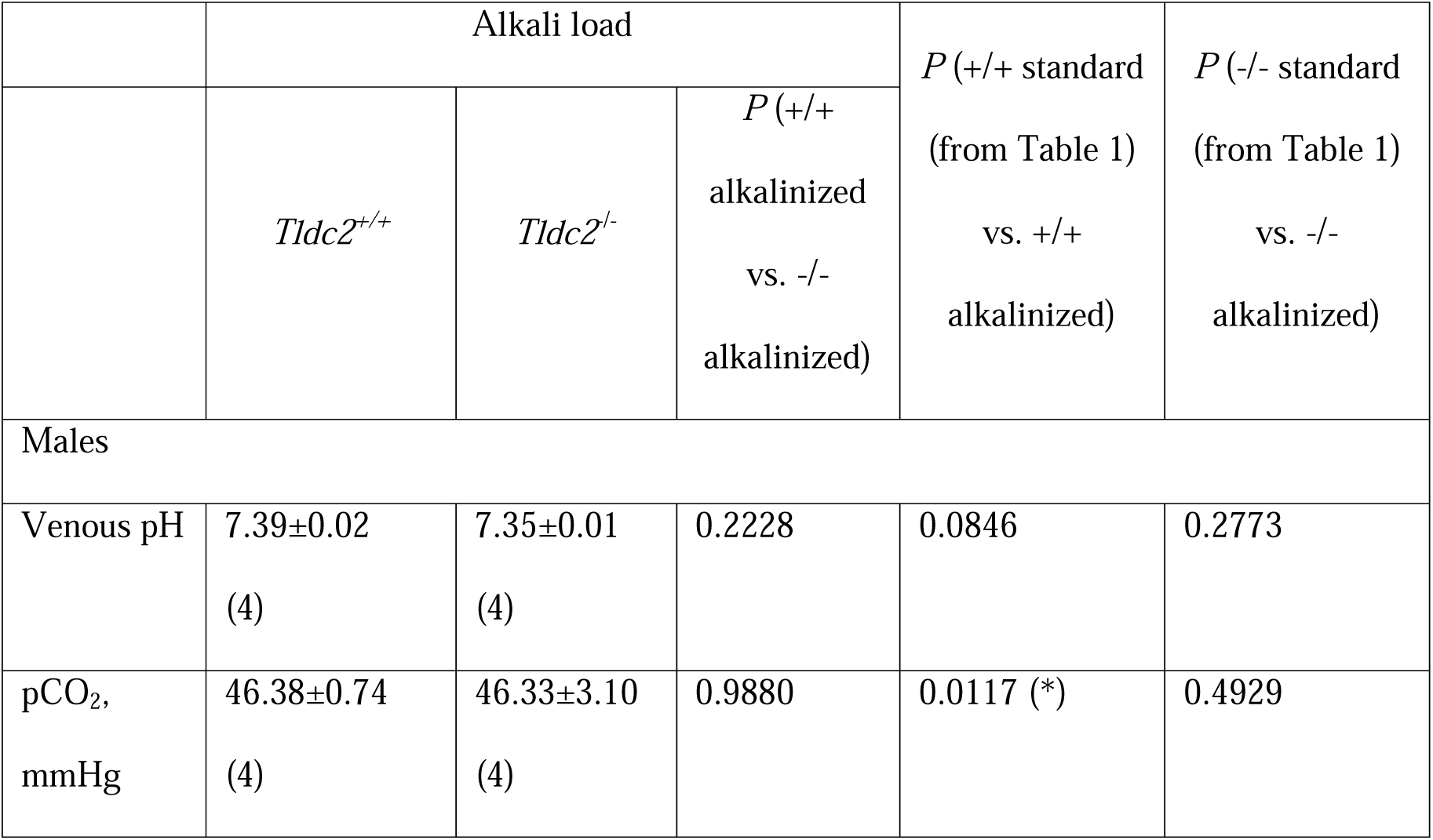
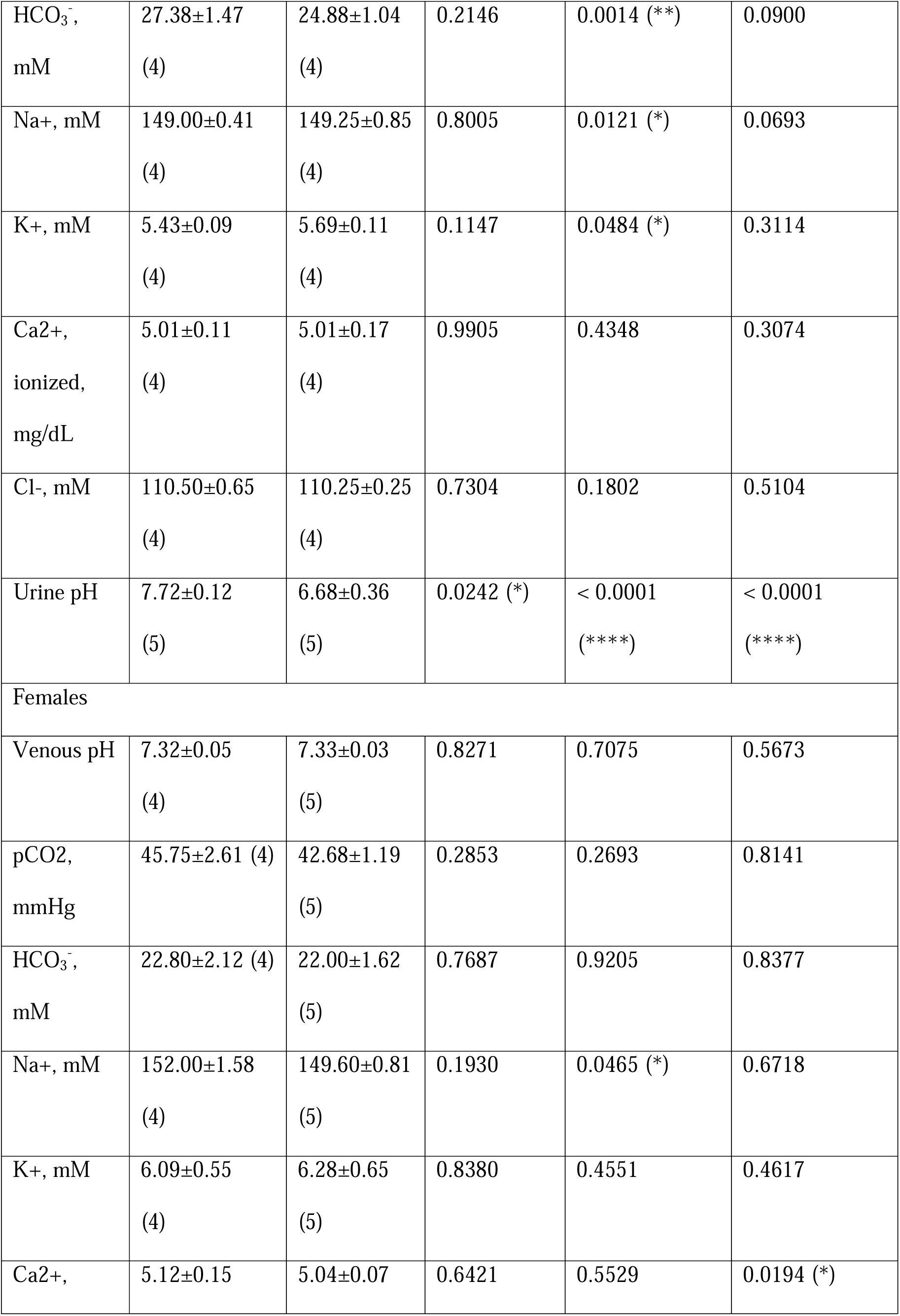
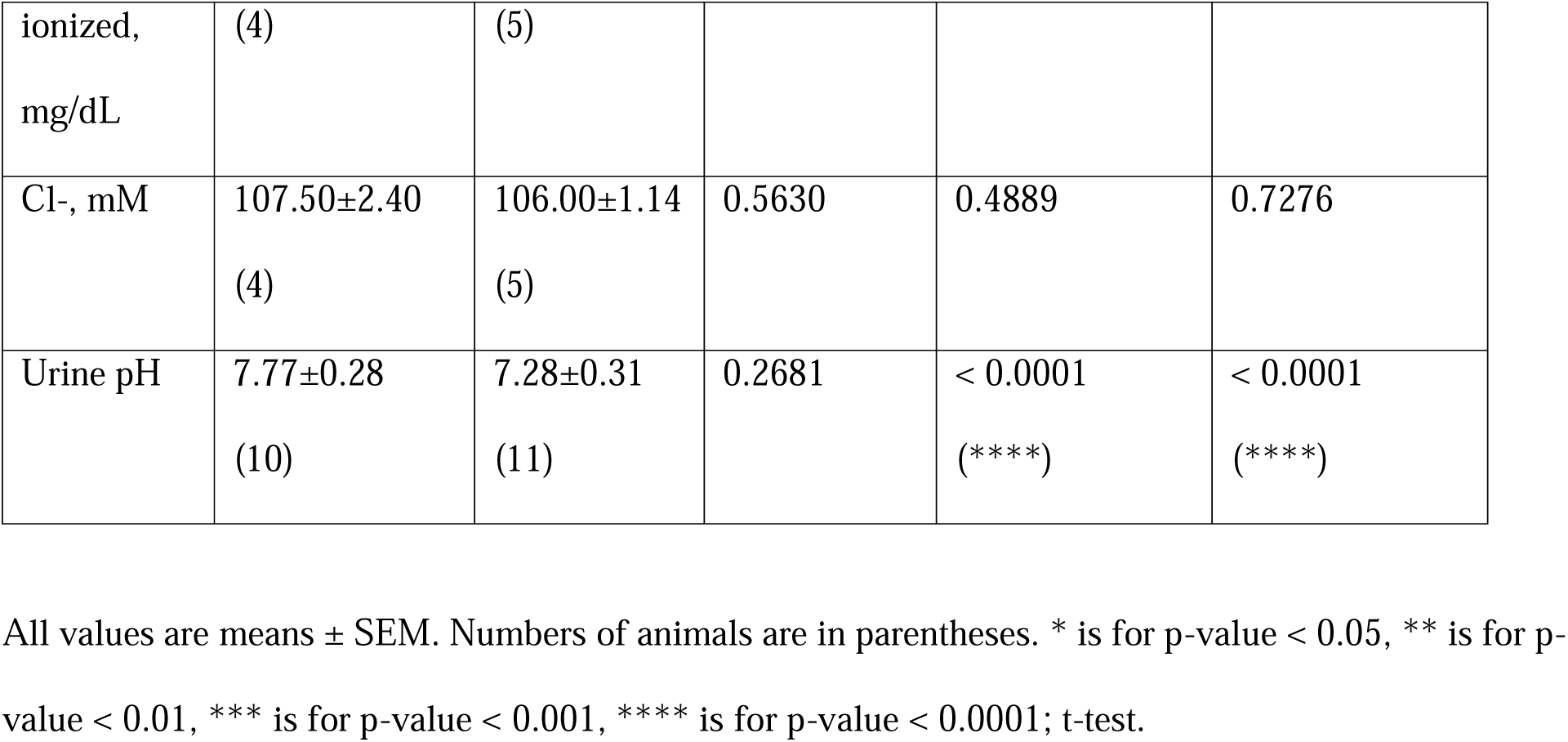
Summary of NaHCO_3_–loading of male and female mice. Venous blood gases, pH and major electrolytes, and urine pH of *Tldc2*^-/-^ knockout and *Tldc2^+/+^* control mice after alkali loading with 0.28 M NaHCO_3_ for 3 days compared to a standard rodent diet.

To study the *Tldc2^-/-^* mice response to both acid and alkali challenge, they were provided with either 0.28 M NH_4_Cl or 0.28 M NaHCO_3_, both for 3 days, in their drinking water. When *Tldc2*^-/-^ and *Tldc2*^+/+^ mice were challenged with 0.28 M NH_4_Cl, urine pH was significantly reduced in *Tldc2*^+/+^ mice as expected (5.78±0.02 vs. 6.13±0.05 in males, Fig. 3 and Table 1; and 5.76±0.07 vs 6.17±0.08 in females; Fig. 4 and Table 1). However, urine pH was not further reduced in *Tldc2*^-/-^ mice apparently because it was already maximally acidic on a standard diet (5.84±0.03 vs. 5.78±0.03 in males, Fig. 3 and Table 1; and 5.77±0.03 vs. 5.88±0.04 in females; Fig. 4 and Table 1). After NH_4_Cl challenge, blood pH was reduced in both *Tldc2*^-/-^ and *Tldc2*^+/+^ male mice but there was no significant difference between genotypes (7.22±0.03 vs. 7.34±0.01 in *Tldc2*^+/+^ males, and 7.21±0.04 vs. 7.33±0.01 in *Tldc2*^-/-^ males, Fig 3 and Table 1). Unlike in male mice, blood pH was significantly reduced only in *Tldc2*^-/-^ and not in *Tldc2*^+/+^ females in response to NH_4_Cl treatment (7.27±0.03 vs 7.35±0.02, Fig. 4 and Table 1).

We then challenged *Tldc2*^-/-^ and *Tldc2*^+/+^ mice with 0.28 M NaHCO_3_, as expected, the urine pH was significantly increased in both *Tldc2*^-/-^ and *Tldc2*^+/+^ male and *Tldc2*^-/-^ and *Tldc2*^+/+^ female mice (Figs. 3 and 4 and Table 2). However, in *Tldc2*^-/-^ males it remained significantly lower than in *Tldc2*^+/+^ control males (6.68±0.36 vs. 7.72±0.12, Fig. 3 and Table 2), suggesting inability of *Tldc2*^-/-^ male mice to efficiently secrete excess bicarbonate into the urine. The *Tldc2*^-/-^ female mice showed a trend toward an impaired ability to excrete HCO3^-^ (7.28±0.31 vs. 7.77±0.28, Fig. 4 and Table 2), but there was a wide spread of urine pH values in both *Tldc2*^+/+^ and *Tldc2*^-/-^ female mice after HCO_3_ ^-^ loading that we found throughout our studies, and that remains unexplained. Then, we found that after NaHCO_3_ challenge *Tldc2*^+/+^ male, but not female, mice developed hyperbicarbonatemia (27.38±1.47 vs. 20.38±0.70 mM in males, Fig. 3 and Table 2; and 22.80±2.12 vs. 22.57±1.19 mM in females; Fig. 4 and Table 2). In comparison with *Tldc2*^+/+^ males, *Tldc2*^-/-^ males were hyperbicarbonatemic even on the standard diet, but after bicarbonate challenge, their blood bicarbonate concentration was not increased further, possibly because an upper limit had been reached (Fig. 3 and Tables 1 and 2). In all these male mice, hyperbicarbonatemia was accompanied by hypercapnia and therefore blood pH stayed within normal range (Fig. 2 and Tables 1 and 2). Thus, neither male nor female *Tldc2*^-/-^ mice developed alkalemia even after bicarbonate challenge.

Finally, we measured concentrations of Na^+^, Cl^-^, Ca^2+^ and K^+^ in blood of *Tldc2*^-/-^ and *Tldc2*^+/+^ both male and female mice, and none of them were significantly different between genotypes under standard, acidotic or alkalotic conditions (Figs. 5 and 6, and Tables 1 and 2), thus *Tldc2* knockout in mice did not significantly affect homeostasis of these electrolytes in blood.

**Figure 5.**
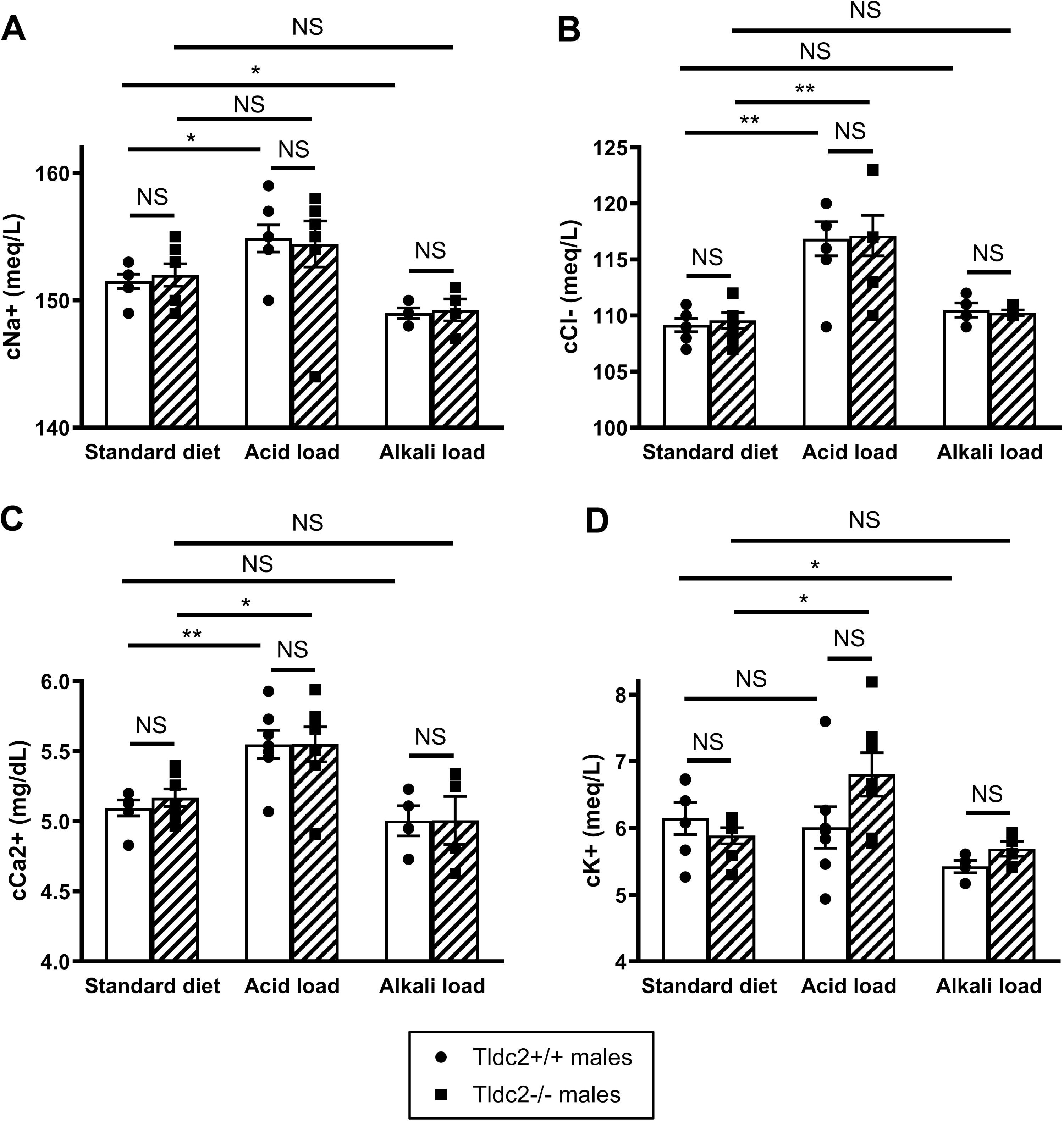
The levels of Na+, Cl-, Ca2+, and K+ in the blood of Tldc2^-/-^ male mice are comparable to those in Tldc2^+/+^ control males when maintained on a standard diet and during acid and alkali challenges. (A) Na+ concentration in *Tldc2^-/-^* male mice is not significantly different from *Tldc2^+/+^* control males on a standard rodent diet. After acid loading with 0.28 M NH_4_Cl for 3 days, Na+ concentration was significantly increased in *Tldc2^+/+^* males only, but it still did not result in significant difference from *Tldc2^-/-^*males. After alkali loading with 0.28 M NaHCO_3_ for 3 days, Na+ concentration was significantly reduced in *Tldc2^+/+^* males only, but it was also not significantly different from *Tldc2^-/-^* males. (B) and (C) Cl- and free Ca2+ concentrations in *Tldc2^-/-^* male mice are not significantly different from Tldc2^+/+^ control males on a standard rodent diet. Both *Tldc2^+/+^* or *Tldc2^-/-^*male mice developed hyperchloremia and hypercalcemia after acid loading with 0.28 M NH_4_Cl for 3 days with no significant difference between genotypes. Both Cl- and free Ca2+ concentrations did not change significantly after alkali loading with 0.28 M NaHCO_3_ for 3 days in ether *Tldc2^+/+^*or *Tldc2^-/-^* males, and there were no statistically significant differences between genotypes. (D) K+ concentration in *Tldc2^-/-^*male mice is not significantly different from Tldc2^+/+^ control males on a standard rodent diet. After acid loading with 0.28 M NH_4_Cl for 3 days, K+ concentration was significantly increased in *Tldc2^-/-^* males only, but it still did not result in significant difference from *Tldc2^+/+^* males. After alkali loading with 0.28 M NaHCO_3_ for 3 days, K+ concentration was significantly reduced in *Tldc2^+/+^* males only, however it did not become significantly different from *Tldc2^-/-^* males. All values are means ± SEM. * is for p-value < 0.05, ** is for p-value < 0.01, t-test. NS = non-significant. The actual p-values and other details can be found in Tables 1 and 2.

**Figure 6.**
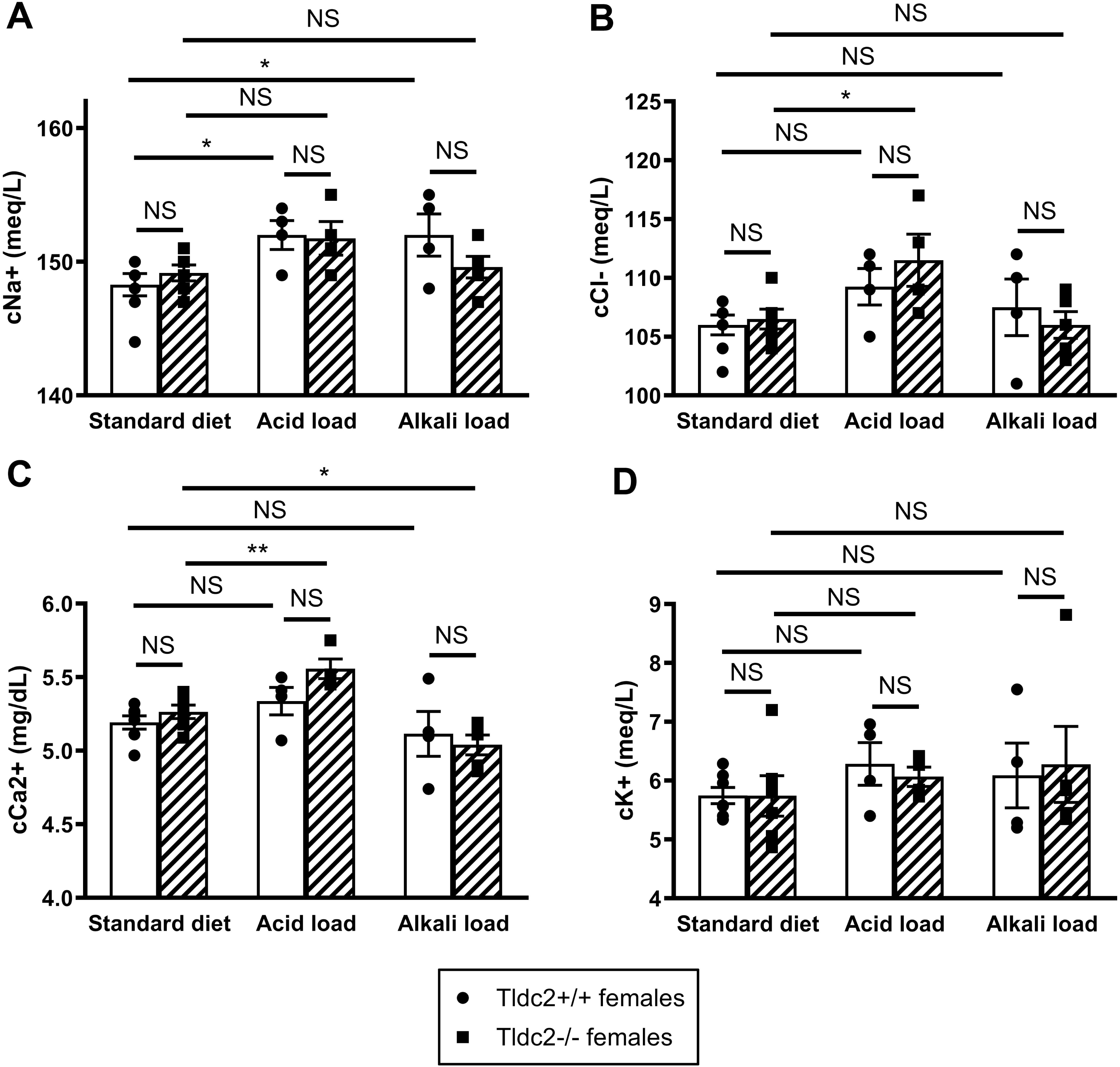
The levels of Na+, Cl-, Ca2+, and K+ in the blood of Tldc2^-/-^ female mice are comparable to those in Tldc2^+/+^ control females when maintained on a standard diet and during acid and alkali challenges. (A) Na+ concentration in *Tldc2^-/-^* female mice is not significantly different from *Tldc2^+/+^* control females on a standard rodent diet. After acid loading with 0.28 M NH_4_Cl for 3 days, Na+ concentration was significantly increased in *Tldc2^+/+^* females only, but it still did not result in significant difference from *Tldc2^-/-^*females. After alkali loading with 0.28 M NaHCO_3_ for 3 days, Na+ concentration was also significantly increased in *Tldc2^+/+^* females only, but it was also not significantly different from *Tldc2^-/-^* females. (B) Cl-concentration in *Tldc2^-/-^*female mice is not significantly different from Tldc2^+/+^ control females on a standard rodent diet. Only *Tldc2^-/-^* female mice developed hyperchloremia after acid loading with 0.28 M NH_4_Cl for 3 days but it did not result in significant difference from *Tldc2^+/+^*females. After alkali loading with 0.28 M NaHCO_3_ for 3 days Cl-concentration did not change significantly in ether *Tldc2^+/+^* or *Tldc2^-/-^* females, and there was no statistically significant difference between genotypes. (C) Free Ca2+ concentration in *Tldc2^-/-^*female mice is not significantly different from Tldc2^+/+^ control females on a standard rodent diet. Only *Tldc2^-/-^* female mice developed hypercalcemia after acid loading with 0.28 M NH_4_Cl for 3 days, and hypocalcemia after alkali loading with 0.28 M NaHCO_3_ for 3 days. Nevertheless, both treatments did not result in significant difference in free Ca2+ concentration between *Tldc2^+/+^* and *Tldc2^-/-^* females. (D) K+ concentration in *Tldc2^-/-^* female mice is not significantly different from Tldc2^+/+^ control females on a standard rodent diet. K+ concentration did not change significantly after acid loading with 0.28 M NH_4_Cl for 3 days or after alkali loading with 0.28 M NaHCO_3_ for 3 days in ether *Tldc2^+/+^* or *Tldc2^-/-^*males and there was no statistically significant difference between genotypes. All values are means ± SEM. * is for p-value < 0.05, ** is for p-value < 0.01, t-test. NS = non-significant. The actual p-values and other details can be found in Tables 1 and 2.

### Tldc2 knockout mice have fewer B-intercalated cells under normal and bicarbonate challenged conditions

To study the molecular and cellular mechanisms underlying the observed aciduria, we first studied the expression of the V-ATPase B1 subunit and pendrin in the kidneys of *Tldc2*^-/-^ mice by Western blotting but did not see any significant difference in their expression in comparison with control *Tldc2^+/+^* mice (Fig. 7). Then, after challenging *Tldc2*^-/-^ and *Tldc2*^+/+^ mice with bicarbonate, we examined the localization of the B1 V-ATPase and pendrin in B-ICs and non-A non-B ICs in the cortex. In untreated *Tldc2*^-/-^ B-ICs, we quantified decreased basolateral localization of the B1 subunit of the V-ATPase relative to control B-ICs, which suggested that Tldc2 may play a role in the delivery or accumulation of B1 into the basolateral membrane in B-ICs (Fig. 8*A* and *B*). However, after treatment with bicarbonate, there was a minor (but not quite significant) trend towards increased basolateral localization of B1 in B-ICs from *Tldc2*^-/-^ mice (p = 0.0526) (Fig. 8*C*) indicting that in the absence of Tldc2 there may be other mechanisms regulating the V-ATPase dependent cellular response to bicarbonate. There was no significant difference in the localization of B1 in non-A non-B cells from *Tldc2*^-/-^and control mice under control or bicarbonate-treated conditions (Fig. 8*D*). We, also, saw no difference in the localization of pendrin in either B-ICs or non-A non-B ICs in untreated or bicarbonate-treated animals (Fig. 8*C* and *D*). Furthermore, to try and understand the marked decrease in B-ICs in *Tldc2*^-/-^ mice, we quantified the percentage of non-A non-B and B-ICs within the total population of pendrin-positive cells, as well as the ratio of pendrin-positive cells (non-A ICs) to total ICs under unchallenged control conditions and after challenging the animals with bicarbonate (Fig. 9). Under baseline conditions, *Tldc2*^-/-^ animals had significantly fewer B-ICs (Fig. 9*A*) and a correspondingly higher percentage of non-A non-B ICs relative to control mice (Fig. 9*B*), suggesting that these populations of cells may interconvert, and this transition may be affected by the loss of Tldc2. This trend was maintained after treatment with bicarbonate with *Tldc2*^-/-^ mice having significantly fewer B-ICs and more non-A non-B ICs than their wildtype counterparts (Fig 9. *A* and *B*). Finally, we examined the ratio of pendrin-positive ICs (non-A ICs) to total ICs under unchallenged and base challenged conditions and found no significant differences (Fig. 9*C*). This suggests that loss of Tldc2 affects only the subpopulation of pendrin-positive non-A non-B and B-ICs, while not altering the population of AE1-positive A-ICs.

**Figure 7.**
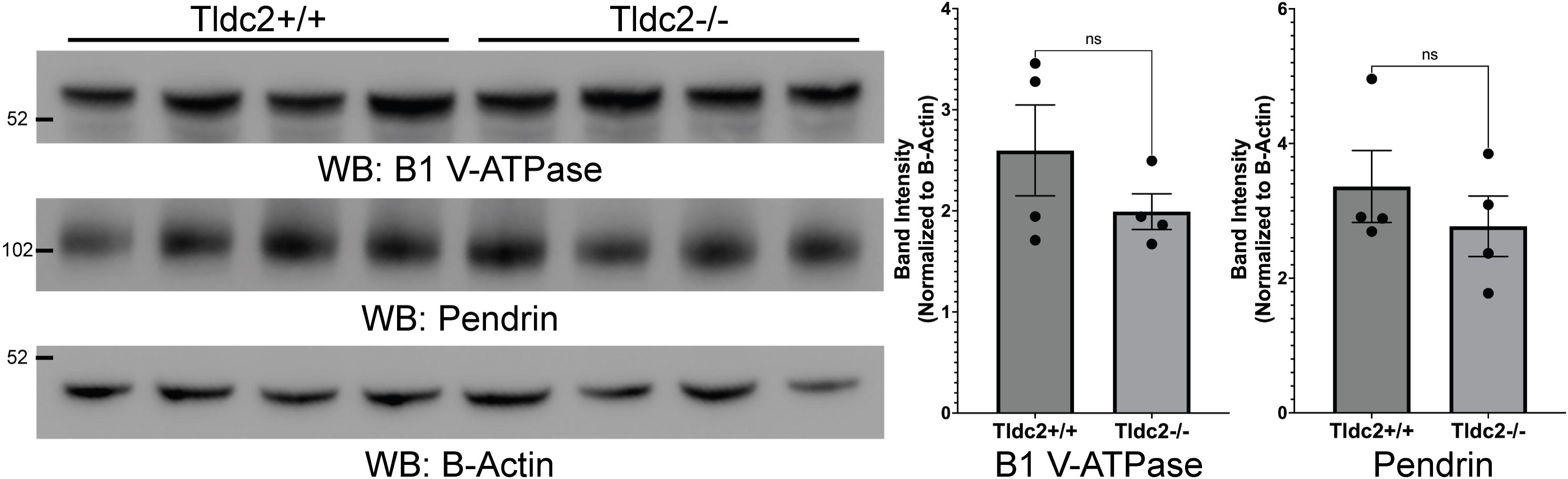
Protein expression of the B1 subunit of the V-ATPase and pendrin are not significantly different in kidneys from *Tldc2^+/+^* and *Tldc2^-/-^*mice. (Left) Western blots of total kidney lysates prepared from wildtype *Tldc2^+/+^* (WT) and knockout *Tldc2^-/-^* (KO) mice probed with specific antibodies against the B1 subunit of the V-ATPase and pendrin, and B-Actin. (Right) Quantification of B1 and pendrin band intensities normalized to B-Actin band intensity shows no difference in expression in total kidney lysates from *Tldc2^+/+^* and *Tldc2^-/-^*mice. ns – non-significant.

**Figure 8.**
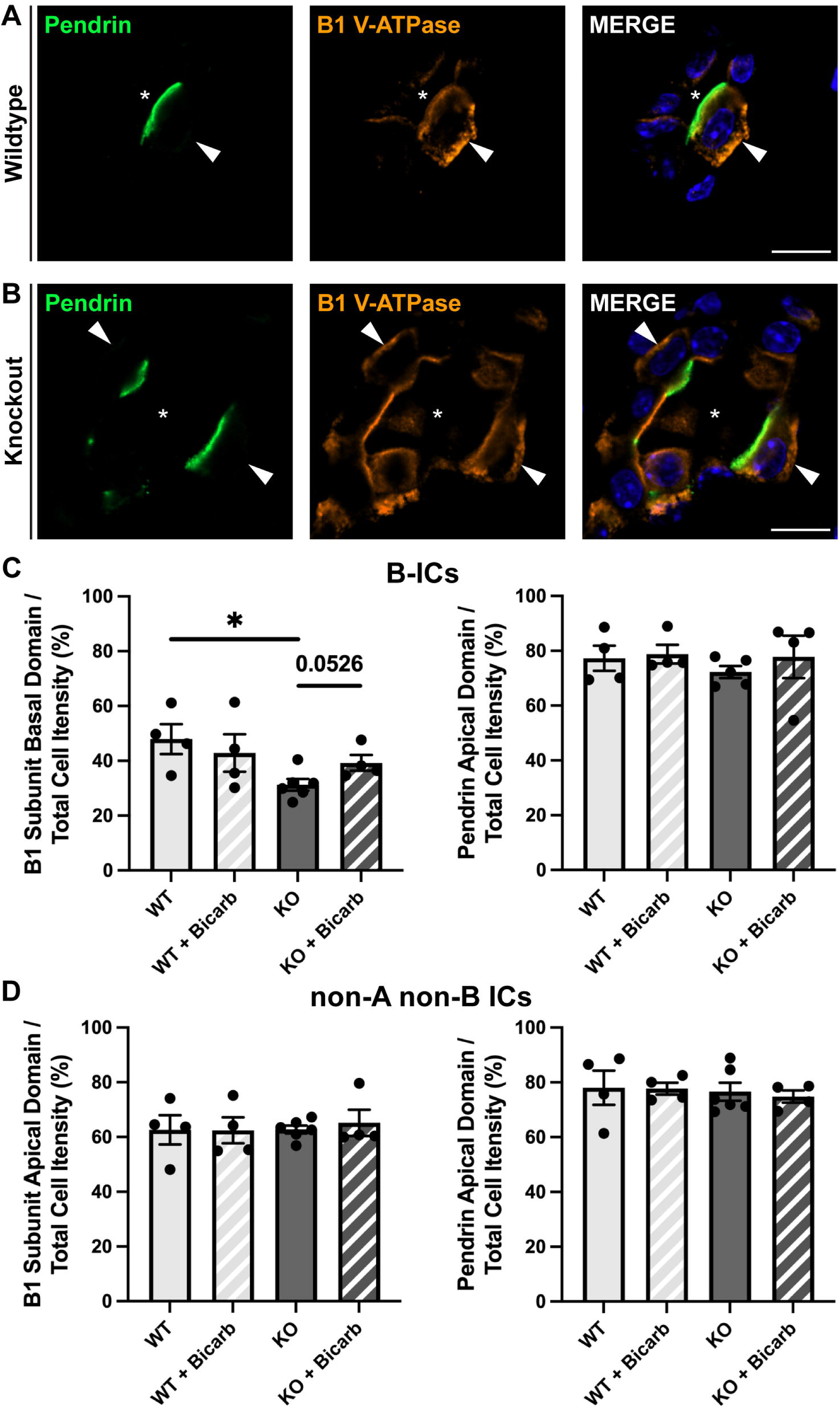
Basal localization of the B1 subunit of the V-ATPase is decreased in B-ICs from *Tldc2^-/-^*mice relative to *Tldc2^+/+^* mice, while apical B1 localization in non-A non-B ICs is not different and apical pendrin localization is unaffected in either cell type. (A-B) Immunofluorescence images taken from the cortex of kidneys from (A) wildtype *Tldc2^+/+^* (WT) and (B) knockout *Tldc2^-/-^*(KO) mice stained with specific antibodies against pendrin (green) and the B1 subunit of the V-ATPase (orange). B1 is basally polarized in B-ICs from *Tldc2^+/+^* mice but shows a more diffuse basolateral distribution in B-ICs from *Tldc2^-/-^* mice. Apical pendrin expression with basolateral B1 expression was used to identify B-ICs (arrowheads). Nuclei are labeled with DAPI (blue). Tubule lumen is indicated with an asterisk (*). Scale bars = 20 µM. **(C)** Quantification of % basal localization of B1 V-ATPase and % apical localization of pendrin relative to total cell localization in B-ICs. There is significantly less basal polarization of B1 in *Tldc2^-/-^* B-ICs compared to *Tldc2^+/+^*B-ICs under untreated conditions, while pendrin localization is not affected. (D) Quantification of % apical localization of the B1 subunit and % apical localization of pendrin relative to total cell localization in non-A non-B ICs showed no significant differences between *Tldc2^-/-^* and *Tldc2^+/+^* mice. Data were analyzed by *t*-test with values reported as means SEM with *P*-values ≤ .05 considered significant, with * denoting a *P*-value ≤ .05.

**Figure 9.**
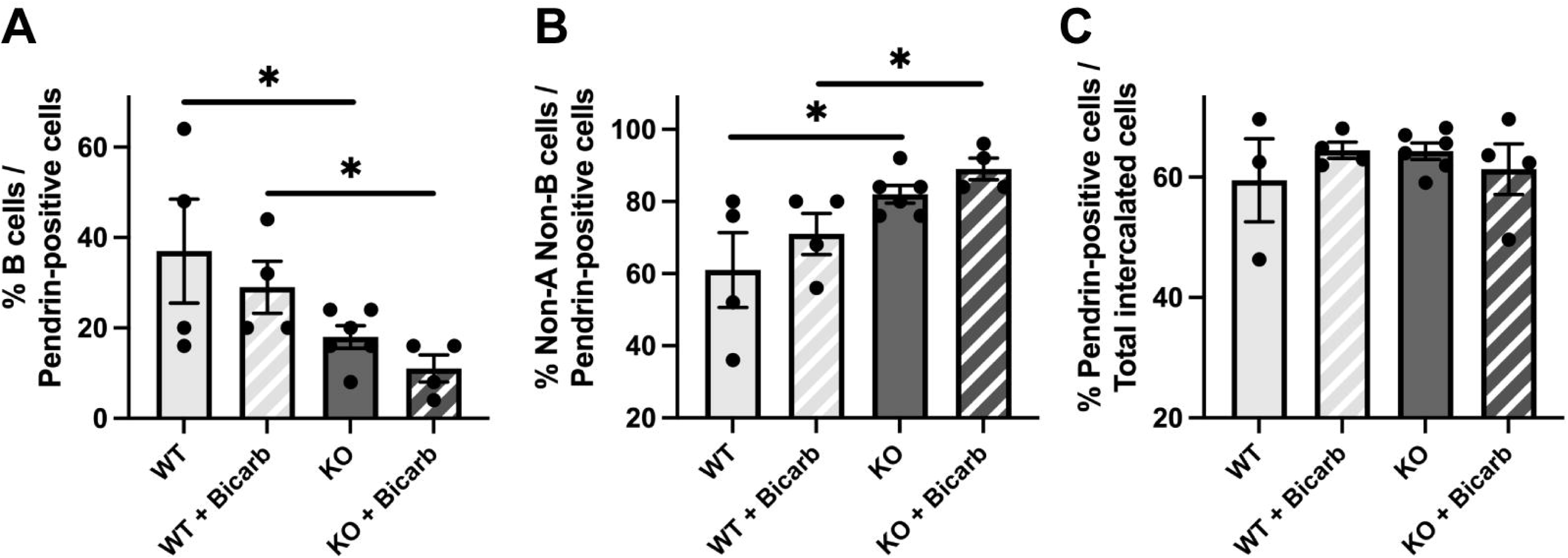
Total number of B-type intercalated cells (B-ICs) is decreased with a corresponding increase in non-A non-B ICs in kidneys from *Tldc2^-/-^* mice, while their sum (all pendrin-positive cells) and the number of A-ICs are not significantly different. (A) Quantification of % B-ICs and (B) % non-A non-B ICs relative to total pendrin-positive cells shows significantly fewer B-ICs with a corresponding increase in non-A non-B ICs in *Tldc2^-/-^* kidneys compared to *Tldc2^+/+^*kidneys under both untreated and bicarbonate-challenged conditions. (C) Quantification of % pendrin-positive cells relative to total ICs in *Tldc2^-/-^*and *Tldc2^+/+^* kidneys shows no significant differences under both untreated and bicarbonate-treated conditions. Data were analyzed by *t*-test with values reported as means SEM with *P*-values ≤ .05 considered significant, with * denoting a *P*-value ≤ .05.

## DISCUSSION

Our recent discovery that TLDc proteins interact with the V-ATPase, and our long-standing interest in V-ATPase function in the kidney prompted us to explore if these proteins regulate the kidney V-ATPase and participate in maintaining acid-base homeostasis in mammals. Initially we studied one of these TLDc proteins, Ncoa7, that is highly expressed in proton-secreting A-ICs (26). We demonstrated that *Ncoa7* knockout mice have some features of incomplete distal renal tubular acidosis (dRTA) and proposed that other TLDc proteins may also regulate V-ATPase function in the kidney in a cell-specific manner based on their expression patterns. Therefore, we investigated here the role of another V-ATPase interacting TLDc protein, Tldc2, in mouse kidney, using a novel *Tldc2* knockout mouse model. First, we examined which cells express Tldc2 in the wild-type kidney. In agreement with previous high-throughput transcriptomics studies, we found that Tldc2 is highly differentially expressed in B-ICs, but in addition we also detected expression using RNAScope in some non-A, non-B ICs, as well. Both of these cell types are pendrin-positive, but while the function of B-ICs in bicarbonate secretion and chloride absorption via pendrin is established (6), the physiological function of non-A, non-B cells is unclear. It is possible that they are partially differentiated precursor cells that give rise to either A-ICs or B-ICs in response to changes in whole-body acid-base balance, or possibly that they are B-ICs in transition between different functional states. In recent years advances in single-cell RNA sequencing have allowed many IC clusters to be distinguished, although whether they are truly functionally unique IC subtypes or, as mentioned above, various transitional states of the originally described A-ICs and B-ICs remains to be determined (2,27). Nonetheless, both B-ICs and non-A non-B ICs express pendrin at their apical membrane domain, while the apical and/or basolateral (and sometimes diffuse) location of the V-ATPase is variable, as seen in the earliest description of V-ATPase localization in these cells (28).

According to previous studies, Tldc2 binds to the V-ATPase (11) and induces its disassembly (12). It has also been shown that V-ATPase disassembly is required prior to the fusion of V-ATPase-containing pre-synaptic vesicles with the plasma membrane and is necessary for its recycling at the neuronal synapse (29). We propose, that in kidney ICs, V-ATPase disassembly is also necessary for its delivery to the plasma membrane where re-assembly then occurs to initiate proton secretion. Thus, by inducing some degree of V-ATPase disassembly in B-ICs under normal conditions, Tldc2 may facilitate V-ATPase localization and recycling at the basolateral membrane of these cells. It is also possible, that Tldc2 is involved in targeting the V-ATPase to the basolateral membrane during the differentiation of B-ICs. Indeed, we found that in Tldc2^-/-^ mice, there is significantly less V-ATPase B1 subunit at the basolateral membrane of B-ICs. In addition, there were significantly fewer classical B-ICs in these mice overall, but significantly more non-A non-B IC. If V-ATPase delivery to the basolateral membrane of pendrin positive ICs is inhibited by the absence of Tldc2, then these cells may not be visualized as bona-fide B-ICs by our definition, and instead could be categorized as non-A non-B ICs due to the reduction in basolateral V-ATPase immunostaining. Which other proteins might be involved in V-ATPase localization and recycling in these highly specialized cells remains to be determined.

We also studied Tldc2 function at the whole-body level using our *Tldc2* global knockout mouse model. Based on the interaction of Tldc2 with the V-ATPase and its highly specific expression in B-ICs in the kidney, we hypothesized that it plays an important role in proton reabsorption and bicarbonate secretion, contributing to acid-base regulation by the kidney. Indeed, we found that urine pH in Tldc2^-/-^ mice was significantly more acidic than in Tldc2^+/+^ mice both with a normal diet and after a NaHCO_3_^-^ challenge, suggesting an inability of these mice to secrete bicarbonate and alkalinize urine as efficiently as Tldc2^+/+^ control mice. However, it did not result in alkalemia and Tldc2^-/-^ mice did not develop alkalosis even during a bicarbonate challenge. In addition, there were no differences in major electrolyte concentrations in blood between Tldc2^-/-^ and Tldc2^+/+^ mice either under standard conditions or after acid/alkali challenges. After bicarbonate challenge Tldc2^-/-^ male, but not Tldc2^-/-^ female mice developed hyperbicarbonatemia, which was partially compensated by hypercapnia due to hypoventilation by the lungs. Overall, Tldc2^-/-^ female mice maintained their acid-base homeostasis more effectively than Tldc2^-/-^ male mice. Sex differences between male and female mice in the renal regulation of acid-base homeostasis, as well as water and electrolyte balance are currently emerging (30–32). Furthermore, there are additional renal mechanisms to prevent hyperbicarbonatemia. For example, it is known that the threshold of the kidney proximal tubule to reabsorb bicarbonate is set around 28 mmol/L (33). Thus, likely because of different compensatory processes, *Tldc2* knockout resulted in a relatively mild renal phenotype in mice. However, our findings clearly highlight the potential role of B-ICs and Tldc2 in protection from alkalosis, in particular, if other (either lung or renal) mechanisms fail. Additionally, persistently low urine pH, as we observed in *Tldc2^-/-^*mice, has other pathophysiological implications. Acidic urine favors uric acid precipitation and may result in kidney stones formation (34). We did not observe stones in kidneys of *Tldc2* knockout mice either on a standard diet or after short-term 3-day NH4Cl acid challenge, but it remains possible that they may form after longer-term challenges in these mice.

Intriguingly, the renal phenotype of *Tldc2* knockout was different from *Atp6v1b1* (B1 subunit of V-ATPase gene) and *Ncoa7* (another member of the TLDc protein family expressed in A-IC) knockout mice, which both had persistently more alkaline urine pH in comparison with controls and demonstrated features of incomplete dRTA (26,35). Of note, *Ncoa7* knockout mice did not develop acidemia after acid load in contrast to *Atp6v1b1* knockout mice, indicating that their knockout phenotype was less severe (26). Additionally, while the protective role of the B1 subunit of V-ATPase against an acid load is well established, only very recently was it shown that it also protects against alkali load (22). In this study, *Atp6v1b1* knockout mice were less protected from metabolic alkalosis than controls and developed more profound hypokalemic alkalosis in response to alkali load (22). We did not observe hypokalemic alkalosis in *Tldc2* knockout mice, but Tldc2^-/-^ male mice did develop hyperbicarbonatemia in response to an alkali load. Thus, interestingly, *Atp6v1b1*, *Ncoa7* and *Tldc2* knockout mice have 3 different phenotypes. While *Atp6v1b1* knockout resulted in both acidosis after acid load and alkalosis after alkali load, *Ncoa7* knockout mice demonstrated only some features of acidosis after acid load, while *Tldc2* knockout mice demonstrated only some features of alkalosis after alkali load. These results are consistent with the different expression patterns of these 3 proteins in the kidney: while the B1 subunit of V-ATPase is expressed in both A-ICs and B-ICs, Ncoa7 is upregulated in A-ICs and Tldc2 is highly specific to B-ICs. This suggests that Ncoa7 and Tldc2 play opposing roles in the kidney.

On one hand, Ncoa7 regulates V-ATPase driven proton secretion in A-ICs, while Tldc2 regulates V-ATPase-dependent proton reabsorption and bicarbonate secretion in B-ICs. It is remarkable, because at the molecular level both Ncoa7 and Tldc2 appear to act in a similar manner: they bind, disassemble, and inhibit V-ATPase activity (12). Despite the similarity of the molecular mechanism, the restricted expression of Ncoa7 and Tldc2 to very particular cell types seems to define their specific roles in the kidney and is important for fine-tuning of V-ATPase function in response to an acid or base challenge, respectively. The phenotype of our *Tldc2* knockout mice also further emphasizes the importance of V-ATPase function in B-ICs and its role in protection from an alkali load.

Although all TLDc proteins interact with V-ATPase, they have very specific, mostly non-overlapping expression patterns in the kidney (27). Therefore, we suggest that the knockout of each of the TLDc proteins in mice will affect kidney function differently and result in a unique phenotype. Thus, we expect that the functional consequences of knocking out the other family members - *Oxr1*, *Tbc1d24* and *Tldc1* - will be different from *Ncoa7* and *Tldc2* knockouts at both the kidney and whole-body level. For example, it is possible that not all of the TLDc proteins are involved in the renal regulation of acid-base homeostasis. We plan to address this question using the same methodology as used for *Ncoa7* and *Tldc2* knockout mice. We are also interested in how the interaction between V-ATPase and TLDc proteins is regulated in the kidney. Indeed, since TLDc binding to V-ATPase results in its inhibition, it should be tightly controlled in a time- and location-specific manner. One of the potential mechanisms of this regulation is by post-translational modifications, such as phosphorylation. Very little is known about that, and this is one of the future directions for V-ATPase research in our laboratory.

## SUPPLEMENTAL MATERIAL

Supplemental Fig. S1.

## ACKNOWLEDGEMENTS

We are grateful to Dr. Susan Wall from Emory University for providing us with the anti-pendrin antibody.

## GRANTS

This work was supported by National Institutes of Health (NIH) Grants R01 DK121848 (to D. B.) and T32 DK007540 (A. F. E.). The Zeiss LSM 800 with Airyscan confocal was purchased using NIH Shared Instrumentation Grant OD021577 (to D. B.). Additional support for the Program in Membrane Biology Microscopy Core comes from the Boston Area Diabetes and Endocrinology Research Center (DK135043) and the Massachusetts General Hospital (MGH) Center for the Study of Inflammatory Bowel Disease (DK043351).

## DISCLOSURES

No conflicts of interest, financial or otherwise, are declared by the authors.

## AUTHOR CONTRIBUTIONS

A.F.E., D.B. and M.M. conceived and designed research; A.F.E., E.C.D., L.J.T. and M.M. performed experiments; A.F.E., E.C.D. and M.M. analyzed data and interpreted results of experiments; M.M. and A.F.E. drafted the manuscript; A.F.E., E.C.D., L.J.T., D.B. and M.M. edited and revised the manuscript; A.F.E., E.C.D., L.J.T., D.B. and M.M. approved the final version of the manuscript.

## Notes

### Competing Interest Statement

The authors have declared no competing interest.

## REFERENCES

1. Eaton, A. F., Merkulova, M., and Brown, D. (2021) The H(+)-ATPase (V-ATPase): from proton pump to signaling complex in health and disease. Am J Physiol Cell Physiol 320, C392–C414

2. Alper, S. L., Natale, J., Gluck, S., Lodish, H. F., and Brown, D. (1989) Subtypes of intercalated cells in rat kidney collecting duct defined by antibodies against erythroid band 3 and renal vacuolar H+-ATPase. Proc Natl Acad Sci U S A 86, 5429–5433

3. Brown, D., Sabolic, I., and Gluck, S. (1992) Polarized targeting of V-ATPase in kidney epithelial cells. J Exp Biol 172, 231–243

4. Royaux, I. E., Wall, S. M., Karniski, L. P., Everett, L. A., Suzuki, K., Knepper, M. A., and Green, E. D. (2001) Pendrin, encoded by the Pendred syndrome gene, resides in the apical region of renal intercalated cells and mediates bicarbonate secretion. Proc Natl Acad Sci U S A 98, 4221–4226

5. Tahaei, E., Pham, T. D., Al-Qusairi, L., Grimm, R., Wall, S. M., and Welling, P. A. (2023) Pendrin regulation is prioritized by anion in high-potassium diets. Am J Physiol Renal Physiol 324, F256–F266

6. Wall, S. M., Verlander, J. W., and Romero, C. A. (2020) The Renal Physiology of Pendrin-Positive Intercalated Cells. Physiol Rev 100, 1119–1147

7. Pham, T. D., Verlander, J. W., Chen, C., Pech, V., Kim, H. I., Kim, Y. H., Weiner, I. D., Milne, G. L., Zent, R., Bock, F., Brown, D., Eaton, A., and Wall, S. M. (2024) Angiotensin II acts through Rac1 to upregulate pendrin: role of NADPH oxidase. Am J Physiol Renal Physiol 326, F202–F218

8. Kim, Y. H., Kwon, T. H., Frische, S., Kim, J., Tisher, C. C., Madsen, K. M., and Nielsen, S. (2002) Immunocytochemical localization of pendrin in intercalated cell subtypes in rat and mouse kidney. Am J Physiol Renal Physiol 283, F744–754

9. Song, H. K., Kim, W. Y., Lee, H. W., Park, E. Y., Han, K. H., Nielsen, S., Madsen, K. M., and Kim, J. (2007) Origin and fate of pendrin-positive intercalated cells in developing mouse kidney. J Am Soc Nephrol 18, 2672–2682

10. Merkulova, M., Paunescu, T. G., Azroyan, A., Marshansky, V., Breton, S., and Brown, D. (2015) Mapping the H(+) (V)-ATPase interactome: identification of proteins involved in trafficking, folding, assembly and phosphorylation. Sci Rep 5, 14827

11. Eaton, A. F., Brown, D., and Merkulova, M. (2021) The evolutionary conserved TLDc domain defines a new class of (H(+))V-ATPase interacting proteins. Sci Rep 11, 22654

12. Oot, R. A., and Wilkens, S. (2024) Human V-ATPase function is positively and negatively regulated by TLDc proteins. Structure 32, 989–1000 e1006

13. Khan, M. M., Lee, S., Couoh-Cardel, S., Oot, R. A., Kim, H., Wilkens, S., and Roh, S. H. (2022) Oxidative stress protein Oxr1 promotes V-ATPase holoenzyme disassembly in catalytic activity-independent manner. EMBO J 41, e109360

14. Klossel, S., Zhu, Y., Amado, L., Bisinski, D. D., Ruta, J., Liu, F., and Gonzalez Montoro, A. (2024) Yeast TLDc domain proteins regulate assembly state and subcellular localization of the V-ATPase. EMBO J 43, 1870–1897

15. Volkert, M. R., Elliott, N. A., and Housman, D. E. (2000) Functional genomics reveals a family of eukaryotic oxidation protection genes. Proc Natl Acad Sci U S A 97, 14530–14535

16. Durand, M., Kolpak, A., Farrell, T., Elliott, N. A., Shao, W., Brown, M., and Volkert, M. R. (2007) The OXR domain defines a conserved family of eukaryotic oxidation resistance proteins. BMC Cell Biol 8, 13

17. Finelli, M. J., Sanchez-Pulido, L., Liu, K. X., Davies, K. E., and Oliver, P. L. (2016) The Evolutionarily Conserved Tre2/Bub2/Cdc16 (TBC), Lysin Motif (LysM), Domain Catalytic (TLDc) Domain Is Neuroprotective against Oxidative Stress. J Biol Chem 291, 2751–2763

18. Wang, J., Rousseau, J., Kim, E., Ehresmann, S., Cheng, Y. T., Duraine, L., Zuo, Z., Park, Y. J., Li-Kroeger, D., Bi, W., Wong, L. J., Rosenfeld, J., Gleeson, J., Faqeih, E., Alkuraya, F. S., Wierenga, K. J., Chen, J., Afenjar, A., Nava, C., Doummar, D., Keren, B., Juusola, J., Grompe, M., Bellen, H. J., and Campeau, P. M. (2019) Loss of Oxidation Resistance 1, OXR1, Is Associated with an Autosomal-Recessive Neurological Disease with Cerebellar Atrophy and Lysosomal Dysfunction. Am J Hum Genet 105, 1237–1253

19. Doan, R. N., Lim, E. T., De Rubeis, S., Betancur, C., Cutler, D. J., Chiocchetti, A. G., Overman, L. M., Soucy, A., Goetze, S., Autism Sequencing, C., Freitag, C. M., Daly, M. J., Walsh, C. A., Buxbaum, J. D., and Yu, T. W. (2019) Recessive gene disruptions in autism spectrum disorder. Nat Genet 51, 1092–1098

20. Mucha, B. E., Hennekam, R. C. M., Sisodiya, S., and Campeau, P. M. (1993) TBC1D24-Related Disorders. in GeneReviews((R)) (Adam, M. P., Feldman, J., Mirzaa, G. M., Pagon, R. A., Wallace, S. E., and Amemiya, A. eds.), Seattle (WA). pp

21. Chen, L., Lee, J. W., Chou, C. L., Nair, A. V., Battistone, M. A., Paunescu, T. G., Merkulova, M., Breton, S., Verlander, J. W., Wall, S. M., Brown, D., Burg, M. B., and Knepper, M. A. (2017) Transcriptomes of major renal collecting duct cell types in mouse identified by single-cell RNA-seq. Proc Natl Acad Sci U S A 114, E9989–E9998

22. Bourgeois, S., Kovacikova, J., Bugarski, M., Bettoni, C., Gehring, N., Hall, A., and Wagner, C. A. (2024) The B1 H + -ATPase (Atp6v1b1) Subunit in Non-Type A Intercalated Cells is Required for Driving Pendrin Activity and the Renal Defense Against Alkalosis. J Am Soc Nephrol 35, 7–21

23. Paunescu, T. G., Ljubojevic, M., Russo, L. M., Winter, C., McLaughlin, M. M., Wagner, C. A., Breton, S., and Brown, D. (2010) cAMP stimulates apical V-ATPase accumulation, microvillar elongation, and proton extrusion in kidney collecting duct A-intercalated cells. Am J Physiol Renal Physiol 298, F643–654

24. Brown, D., Lydon, J., McLaughlin, M., Stuart-Tilley, A., Tyszkowski, R., and Alper, S. (1996) Antigen retrieval in cryostat tissue sections and cultured cells by treatment with sodium dodecyl sulfate (SDS). Histochem Cell Biol 105, 261–267

25. Schindelin, J., Arganda-Carreras, I., Frise, E., Kaynig, V., Longair, M., Pietzsch, T., Preibisch, S., Rueden, C., Saalfeld, S., Schmid, B., Tinevez, J. Y., White, D. J., Hartenstein, V., Eliceiri, K., Tomancak, P., and Cardona, A. (2012) Fiji: an open-source platform for biological-image analysis. Nat Methods 9, 676-682

26. Merkulova, M., Paunescu, T. G., Nair, A. V., Wang, C. Y., Capen, D. E., Oliver, P. L., Breton, S., and Brown, D. (2018) Targeted deletion of the Ncoa7 gene results in incomplete distal renal tubular acidosis in mice. Am J Physiol Renal Physiol 315, F173–F185

27. Ransick, A., Lindstrom, N. O., Liu, J., Zhu, Q., Guo, J. J., Alvarado, G. F., Kim, A. D., Black, H. G., Kim, J., and McMahon, A. P. (2019) Single-Cell Profiling Reveals Sex, Lineage, and Regional Diversity in the Mouse Kidney. Dev Cell 51, 399–413 e397

28. Brown, D., Hirsch, S., and Gluck, S. (1988) An H+-ATPase in opposite plasma membrane domains in kidney epithelial cell subpopulations. Nature 331, 622–624

29. Bodzeta, A., Kahms, M., and Klingauf, J. (2017) The Presynaptic v-ATPase Reversibly Disassembles and Thereby Modulates Exocytosis but Is Not Part of the Fusion Machinery. Cell Rep 20, 1348–1359

30. Eaton, A. F., Danielson, E. C., Capen, D., Merkulova, M., and Brown, D. (2024) Dmxl1 Is an Essential Mammalian Gene that Is Required for V-ATPase Assembly and Function In Vivo. Function (Oxf) 5

31. Nair, A. V., Yanhong, W., Paunescu, T. G., Bouley, R., and Brown, D. (2019) Sex-dependent differences in water homeostasis in wild-type and V-ATPase B1-subunit deficient mice. PLoS One 14, e0219940

32. McDonough, A. A., Harris, A. N., Xiong, L. I., and Layton, A. T. (2024) Sex differences in renal transporters: assessment and functional consequences. Nat Rev Nephrol 20, 21–36

33. Rodriguez Soriano, J., Boichis, H., Stark, H., and Edelmann, C. M., Jr. (1967) Proximal renal tubular acidosis. A defect in bicarbonate reabsorption with normal urinary acidification. Pediatr Res 1, 81–98

34. Song, L., and Maalouf, N. M. (2000) Nephrolithiasis. in Endotext (Feingold, K. R., Anawalt, B., Blackman, M. R., Boyce, A., Chrousos, G., Corpas, E., de Herder, W. W., Dhatariya, K., Dungan, K., Hofland, J., Kalra, S., Kaltsas, G., Kapoor, N., Koch, C., Kopp, P., Korbonits, M., Kovacs, C. S., Kuohung, W., Laferrere, B., Levy, M., McGee, E. A., McLachlan, R., New, M., Purnell, J., Sahay, R., Shah, A. S., Singer, F., Sperling, M. A., Stratakis, C. A., Trence, D. L., and Wilson, D. P. eds.), South Dartmouth (MA). pp

35. Finberg, K. E., Wagner, C. A., Bailey, M. A., Paunescu, T. G., Breton, S., Brown, D., Giebisch, G., Geibel, J. P., and Lifton, R. P. (2005) The B1-subunit of the H(+) ATPase is required for maximal urinary acidification. Proc Natl Acad Sci U S A 102, 13616–13621

