## Supplemental Fig S1 for "Knockout of the V-ATPase interacting protein Tldc2 in B-type kidney intercalated cells impairs urine alkalinization"

**Supplemental Figure S1. Confirmation of RNA scope assay feasibility.**

Immunofluorescence images taken from the cortex of kidneys from wildtype Tldc2$^{+/+}$ mice after performing RNA scope using specific probes designed against **(A)** PPIB, a ubiquitously expressed cell cycle protein, as a positive control and **(B)** DAPB, a bacterial protein that should not be expressed in mammalian tissue, as a negative control. A positive RNA scope signal appears as a single intracellular punctum (orange). Numerous intracellular puncta present after incubation with PPIB (top) indicates that the RNA scope assay works with our preparation. Absence of puncta after incubation with DAPB (bottom) indicates that the RNA scope assay shows no non-specific background activity. Nuclei are labeled with DAPI (blue). Green is autofluorescence. Scale bars = 20 µM.

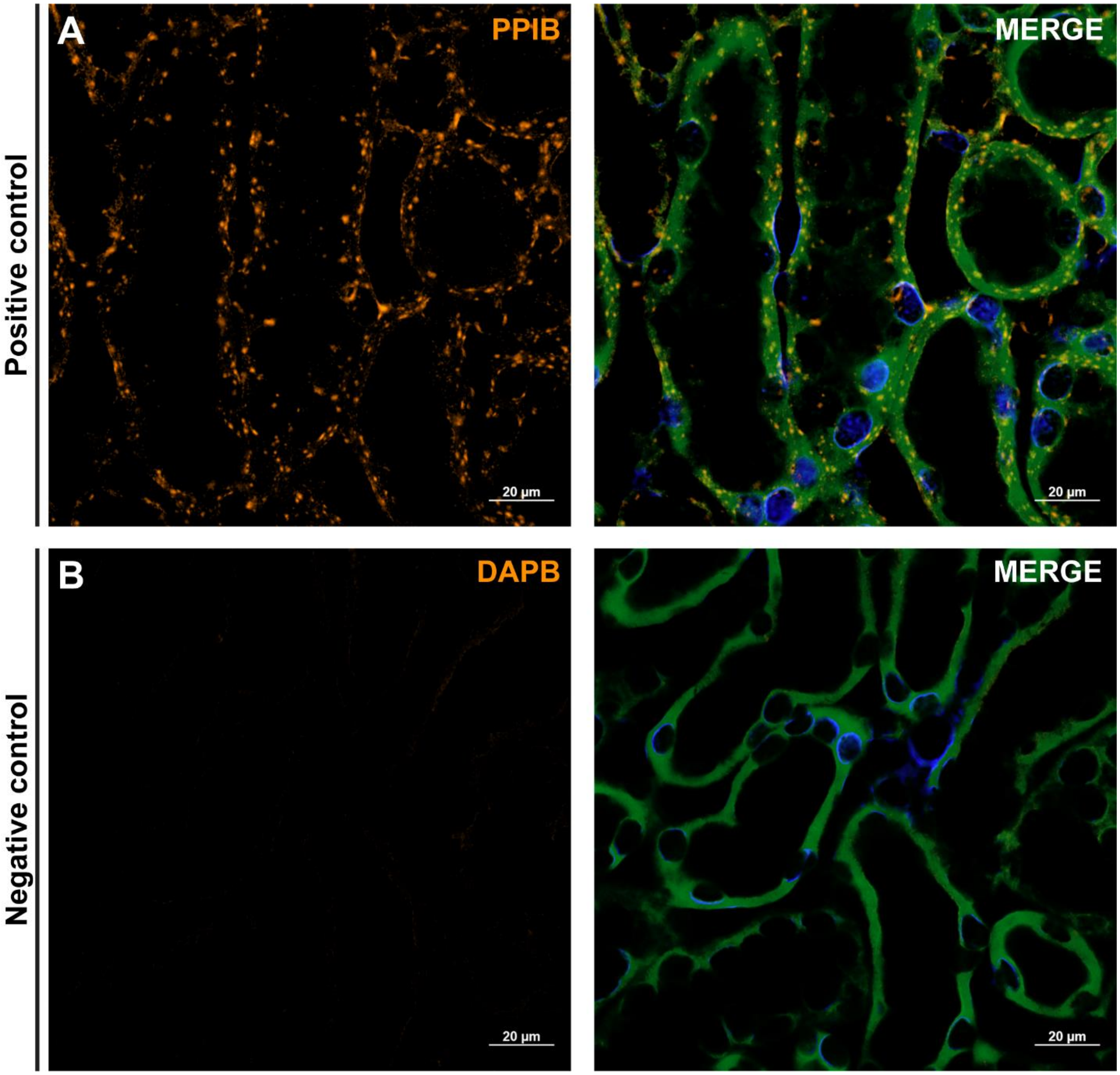
Positive control
A
PPIB
MERGE
20 µm
20 µm
Negative control
B
DAPB
MERGE
20 µm
20 µm
